# The bacterial ESCRT-III PspA rods thin lipid tubules and increase membrane curvature through helix α0 interactions

**DOI:** 10.1101/2025.06.13.659447

**Authors:** Esther Hudina, Stephan Schott-Verdugo, Benedikt Junglas, Mirka Kutzner, Ilona Ritter, Nadja Hellmann, Dirk Schneider, Holger Gohlke, Carsten Sachse

## Abstract

The phage shock protein A (PspA), a bacterial member of the ESCRT-III superfamily, forms rod-shaped helical assemblies that internalize membrane tubules. The N-terminal helix α0 of PspA (and other ESCRT-III members) has been suggested to act as a membrane anchor, the detailed mechanism, however, of how it binds to membranes and eventually triggers membrane fusion and/or fission events remains unclear. By solving a total of 15 cryo-electron microscopy (cryo-EM) structures of PspA and a truncation lacking the N-terminal helix α0 in the presence of EPL membranes, we show in molecular detail how PspA interacts with and remodels membranes: binding of the N-terminal helix α0 in the outer tubular membrane leaflet induces membrane curvature, supporting membrane tubulation by PspA. Detailed molecular dynamics simulations and free energy computations of interactions between the helix α0 and negatively charged membranes suggest a compensating mechanism between helix/membrane interactions and the energy contributions required for membrane bending. The energetic considerations are in line with the membrane structures observed in the cryo-EM images of tubulated membrane vesicles, fragmented vesicles inside tapered PspA rods, and shedded vesicles emerging at the thinner PspA rod ends. Our results provide insights into the molecular determinants and a potential mechanism of vesicular membrane remodeling mediated by a member of the ESCRT-III superfamily.

**Summary:** The rods of bacterial ESCRT-III protein PspA internalize and thin membrane tubules by overcoming the required bending energy through progressive membrane binding of the N-terminal helix in the PspA assembly.

## Introduction

An intact inner membrane is essential for bacterial cell viability, but stressors such as temperature, osmolarity, organic solvents (*e.g.,* ethanol) or phage infections can destabilize the integrity of membranes. To protect the inner membrane, many bacteria activate the bacterial phage shock protein (Psp) response (Bergler et al., 1994; Brissette et al., 1990; Kleerebezem and Tommassen, 1993; Kobayashi et al., 1998). So far, the Psp system is best studied in *E. coli* where the components of the Psp system are encoded by the *pspF-pspABCDE* operon. Here, the proteins PspF, PspA, PspB and PspC form the Psp core elements (Darwin, 2005; Jovanovic et al., 1996; Lloyd et al., 2004). Components of the Psp system have also been identified in other bacteria, cyanobacteria, archaea and chloroplasts (Manganelli and Gennaro, 2017), although strict conservation only exists for PspA and PspC (Huvet et al., 2011; Manganelli and Gennaro, 2017; Popp et al., 2022). However, the Psp-network architecture appears to be more complex than previously thought, showing a large unexpected diversity in the distribution and occurrence of Psp components across bacterial and archaeal species (Popp et al., 2022).

The main effector of the Psp system is PspA, a 25 kDa protein consisting of six α-helices connected by short loops and an elongated hairpin, where the N-terminal helix α0 of the protein remains unfolded in the absence of membranes (Junglas et al., 2021; Osadnik et al., 2015). Proteins of the PspA family (i.e. PspA, Vipp1 and LiaH) form MDa-sized homo-oligomeric assemblies such as carpets, rings and rods with helical symmetry (Gupta et al., 2021; Junglas et al., 2021, 2020; Liu et al., 2021; Standar et al., 2008; Wolf et al., 2010). Interestingly, PspA is related to eukaryotic and archaeal ESCRT-III proteins and has a similar structure (Di Giulio, 2021; Junglas et al., 2021; Liu et al., 2021; Melnikov et al., 2025). Thus, proteins of the PspA family are considered bacterial ESCRT-III proteins. Like their bacterial counterparts, eukaryotic ESCRT-III proteins form homo and hetero-oligomeric assemblies including sheets, strings, rings, filaments, tubules, domes, coils and spirals (Ghazi-Tabatabai et al., 2008; Huber et al., 2020; Lata et al., 2008; McCullough et al., 2015). Although the polymeric assemblies of (bacterial, archaeal and eukaryotic) ESCRT-III proteins come in very different shapes, their monomer structures share the same architecture, and the assemblies also share a similar motif of α5 being packed perpendicularly against the hairpin of α1+2 (Junglas et al., 2021; Schlösser et al., 2023). Also common to all ESCRT-III proteins is their association with membrane remodeling processes. The ESCRT system in eukaryotes assumes critical roles in many cellular processes, including nuclear envelope sealing (Vietri et al., 2015), plasma membrane repair, lysosomal protein degradation (Zhu et al., 2017), retroviral budding and the multivesicular body (MVB) pathway (Schmidt and Teis, 2012). While the basic topology of the budding process is directed away from the cytosol (Hurley and Hanson, 2010), the involved membrane geometries can differ. In the first case, ESCRT-III polymers are required to assemble inside of membrane neck structures to negatively curved membranes as observed *in vitro* and *in vivo* (Azad et al., 2023; Bodon et al., 2011; Hanson et al., 2008). In the second case, ESCRT-III proteins were found to bind to the outside of a membrane tube of positive membrane curvature (Bertin et al., 2020; Huber et al., 2020; McCullough et al., 2015; Moser von Filseck et al., 2020).

How PspA is involved in maintaining the bacterial membrane is so far only poorly understood. It has been shown that PspA binds to negatively charged membranes by the amphipathic N-terminal helix α0, and is capable of membrane remodeling by fusion and fission processes (Junglas et al., 2021; McDonald et al., 2017, 2015). Like Vipp1, a close relative of PspA found in cyanobacteria and chloroplasts, PspA has been suggested to passively protect membranes by forming a protective carpet on the membrane that reduces proton leakage, and/or performing active membrane repair by excising damaged membrane areas and potentially sealing them by fusion with intact membranes (Junglas et al., 2021, 2020; Kobayashi et al., 2007; Siebenaller et al., 2019). However, the molecular details of these processes remained unclear.

We have analyzed the details of PspA-membrane interactions of the thus far unresolved N-terminal helix α0 of PspA of the cyanobacterium *Synechocystis* sp. PCC6803 (form here on PspA). By solving a total of 15 PspA rod cryo-EM structures in the presence of *Escherichia coli* polar lipid (EPL) membranes combined with biophysical and computational methods, we elucidated interactions between PspA and membranes on a molecular level and show that the N-terminal helix α0, located in the lumen of PspA rods, is critical for membrane tubulation: By inserting helix α0 partly into the outer membrane leaflet, PspA generates highly curved EPL membrane tubules. Although the PspA mutant lacking helix α0 shows no structural alterations in the helical rod structure, it can no longer tubulate EPL membranes. Molecular dynamics (MD) simulations and free energy computations suggest a potential orientation of helix α0 on the membrane surface, reveal essential amino acid residues of the N-terminal helix involved in the membrane interaction and provide insights into a compensating mechanism between helix/membrane binding versus the energy contributions required for membrane bending. In the cyro-EM strctures of PspA rods with internalized membranes multiple vesicular structures are visible within a single rod, illustrating a potential membrane remodeling pathway. These results indicate how PspA rods assemble on the membrane surface and, through molecular interactions, can tubulate and thin a lipid bilayer within the rod’s lumen until shedded vesicles emerge.

## Results

### Biophysical characterization of PspA-helix α0 interactions with membranes

To date, the mechanism of PspA interaction with membranes has not been fully characterized as the full-length PspA cryo-EM structure did not contain density for helix α0 (Junglas et al., 2021). Previous studies suggested that the isolated N-terminal peptide α0 adopts an α-helix structure in the presence of negatively charged lipids (Jovanovic et al., 2014; McDonald et al., 2015). To understand the role of helix α0 in membrane interaction and to determine the amino acids relevant for binding to negatively charged lipids, we tested the isolated PspA helix α0 peptide (residues 1-22) and a R6A and R9A (PspA α0 RRAA) mutant with replaced positively charged residues (**Fig. 1A, Suppl. Fig. 1**) and analyzed changes in secondary structure using **c**ircular dichroism (CD) spectroscopy. Upon increasing the EPL concentration, wild-type (WT) α0 revealed a significant reduction in the signal intensity at 220 nm showing formation of α-helical structure induced by membrane-adhesion of the peptide (**Fig. 1B**). Based on the CD data, the α-helix content of the PspA WT α0 peptide was estimated to be ∼52 % at high EPL concentrations, while it was barely present (3.2 %) in the PspA-α0-RRAA sample. The signal intensity at 220 nm did not change in the case of the PspA-α0-RRAA peptide when membranes were added, indicating that its secondary structure is not changing, *i.e.* the petide does not bind to membranes. To investigate whether PspA-α0-RRAA can still bind to membrane surfaces, we additionally monitored changes in the Trp fluorescence in the presence of EPL membranes.

**Figure 1:**
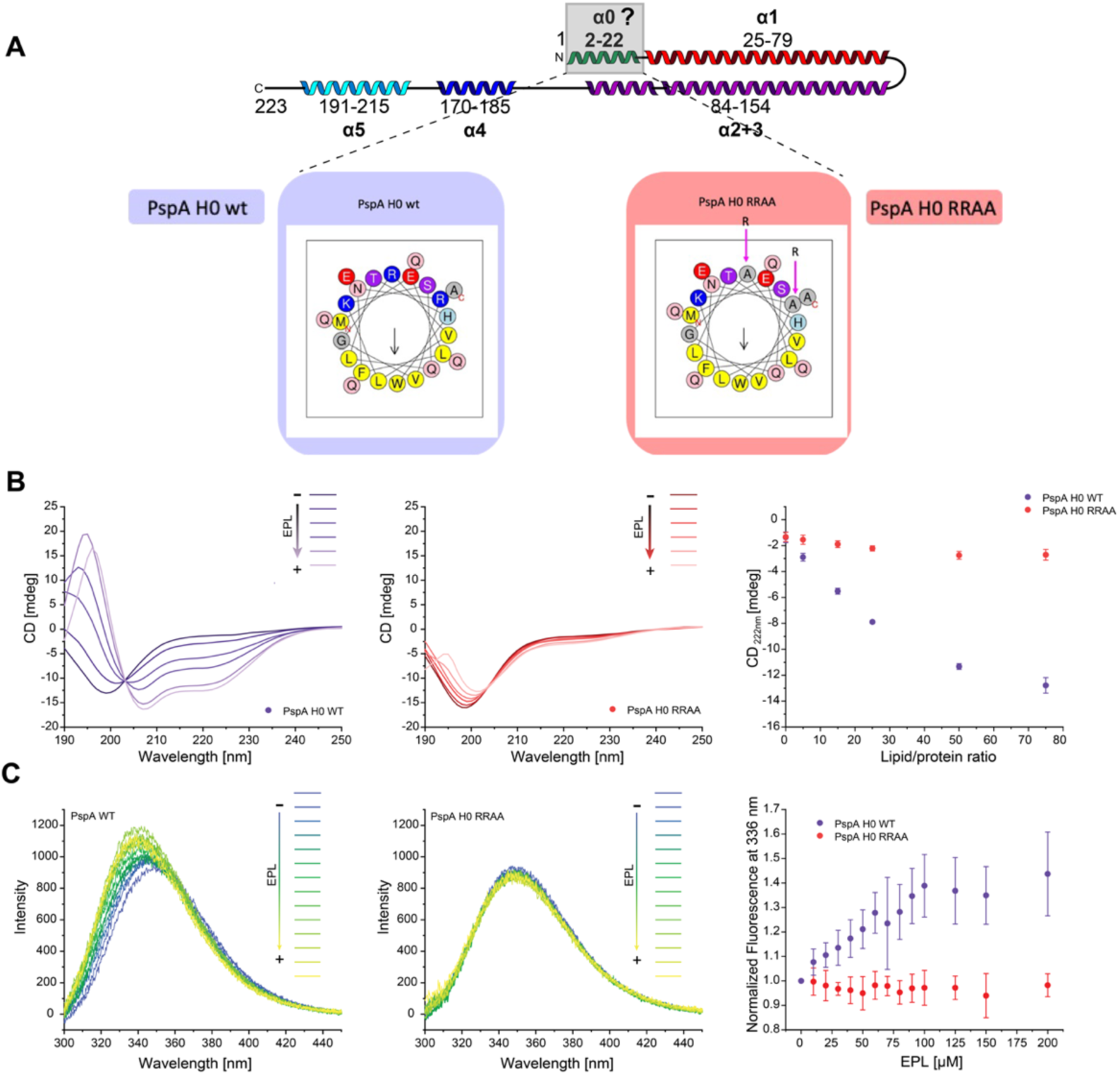
Interaction of PspA’s isolated helix α0 with membranes. **A**: Top: Secondary structure topology plot of the PspA ESCRT-III fold (*α*0 green with grey focus box, *α*1 red, *α*2+3 violet, *α*4 blue, *α*5 cyan). Bottom: Schiffer–Edmundson helix wheel projection of PspA (1-22) wild type (left) and PspA α0 RRAA (right), yellow hydrophobic residues, blue and red polar charged residues, light blue and red polar residues, grey neutral. **B:** Circular dichroism (CD) spectra of helix *α*0 in the presence of increasing EPL concentration (340 µM, 510 µM, 850 µM, 1700 µM, and 2550 µM EPL, left: PspA wild-type (WT) in purple, center: PspA α0 RRAA mutant) in red indicate a membrane-induced increase in α-helix content in the WT depending on the lipid:protein ratio (right). **C**: Changes in the Trp fluorescence emission spectrum in the presence of EPL membranes. Left: PspA WT (purple), center: PspA α0 RRAA (red), right: fluorescence intensity at 336 nm as a function of lipid concentration.

The α0 peptide contains a single Trp residue located in the hydrophobic face of the α-helix projection (see **Fig. 1A**), which likely inserts into the membrane upon helix α0 membrane adhesion. Notably, only the WT, but not the PspA-α0-RRAA peptide, displayed a shift in the Trp fluorescence signal when EPL membranes were added, again indicating that solely the WT-α0-PspA peptide interacts with EPL membranes, whereas the α0 RRAA mutant did not (**Fig. 1C**). Based on the binding curves, the free energy of binding of the WT peptide was estimated to be between Δ*G*^0^ = -3.9 and -3.7 kcal mol^-1^ and between Δ*G*^0^ = -5.7 and -5.3 kcal mol^-1^ for CD spectroscopy and Trp fluorescence, respectively (**Suppl. Fig. 1**). In conclusion, our results suggest that the isolated PspA-α0 peptide binds to membrane surfaces, which involves the concomitant formation of an α-helical structure. The positively charged residues R6 and R9 are critical for helix α0 binding to negatively charged membrane surfaces.

### Full-atom simulation of helix α0 with a lipid bilayer

To investigate at the atomistic level how the PspA-α0 peptide interacts with the lipid membrane, we performed MD simulations of the peptide interacting with a DOPE:DOPG (3:1) membrane. Although initially located 25 Å above the membrane surface (**Fig. 2A**), the peptide bound to the membrane surface in less than 100 ns of simulation time in all 12 replicas while adopting a mainly α-helical secondary structure (**Fig. 2B**, **Suppl. Fig. 1A**). The peptide formed contacts with the membrane through M1, R6, R9 and, less pronounced, K12 (**Fig. 2C**), in agreement with the experimental results supporting the critical role of the positively charged residues R6 and R9 for binding the negatively charged headgroups of the membrane surface. After 2 µs of unbiased MD simulations, the bound peptide was pulled away from the membrane along the membrane normal using adaptive steered MD simulations, until the peptide reached an unbound state with a distance of ∼50 Å from the membrane center. That way, starting structures for umbrella sampling (US) simulations were generated and a free energy profile (potential of mean force (PMF)) of peptide (un)binding was computed. Initially, the PMF indicated strong binding of peptide α0 to the membrane surface (at the water-membrane interface ∼20 Å). Upon unbinding the energy rises steadily until it reaches a plateau beyond 45 Å to the membrane center (**Fig. 2D**). At the lowest energy, the related US structures have a continuous α-helix conformation, with an orientation at the membrane surface constrained by the membrane interactions. With increasing energy, α0 splits up into two α-helical segments introducing a kink at G8 and, at the highest energy, a flexible loop (**Suppl. Fig. 1B-F**). Integration of the PMF shows that the major energetic contribution occurs at a distance to the membrane center below 30 Å (**Suppl. Fig. 1G**). The PMF at distances of 30 to 35 Å to the membrane center marks the detachment of the C-terminal region of the peptide, while the N-terminal region maintains intramolecular hydrogen bonds between E2 and R6/R9, which stabilizes α-helix and kink topology (**Suppl. Fig. 1H**) (Ghosh and Fernández, 2020). The integration of the PMF, considering the loss of configurational and conformational entropy upon membrane-binding of the peptide (see Materials and Methods), yields a binding free energy of Δ*G*^0^ = -6.32 ± 2.07 kcal mol^-1^ for binding of the isolated helix α0 to the model membrane, which is in agreement with the experimentally determined Δ*G*^0^ values. Together, the MD peptide simulations support the experimental results obtained with the isolated helix suggesting binding of the PspA N-terminus to a DOPG:DOPE membrane by forming an amphipathic α-helix next to the lipid headgroups.

**Figure 2:**
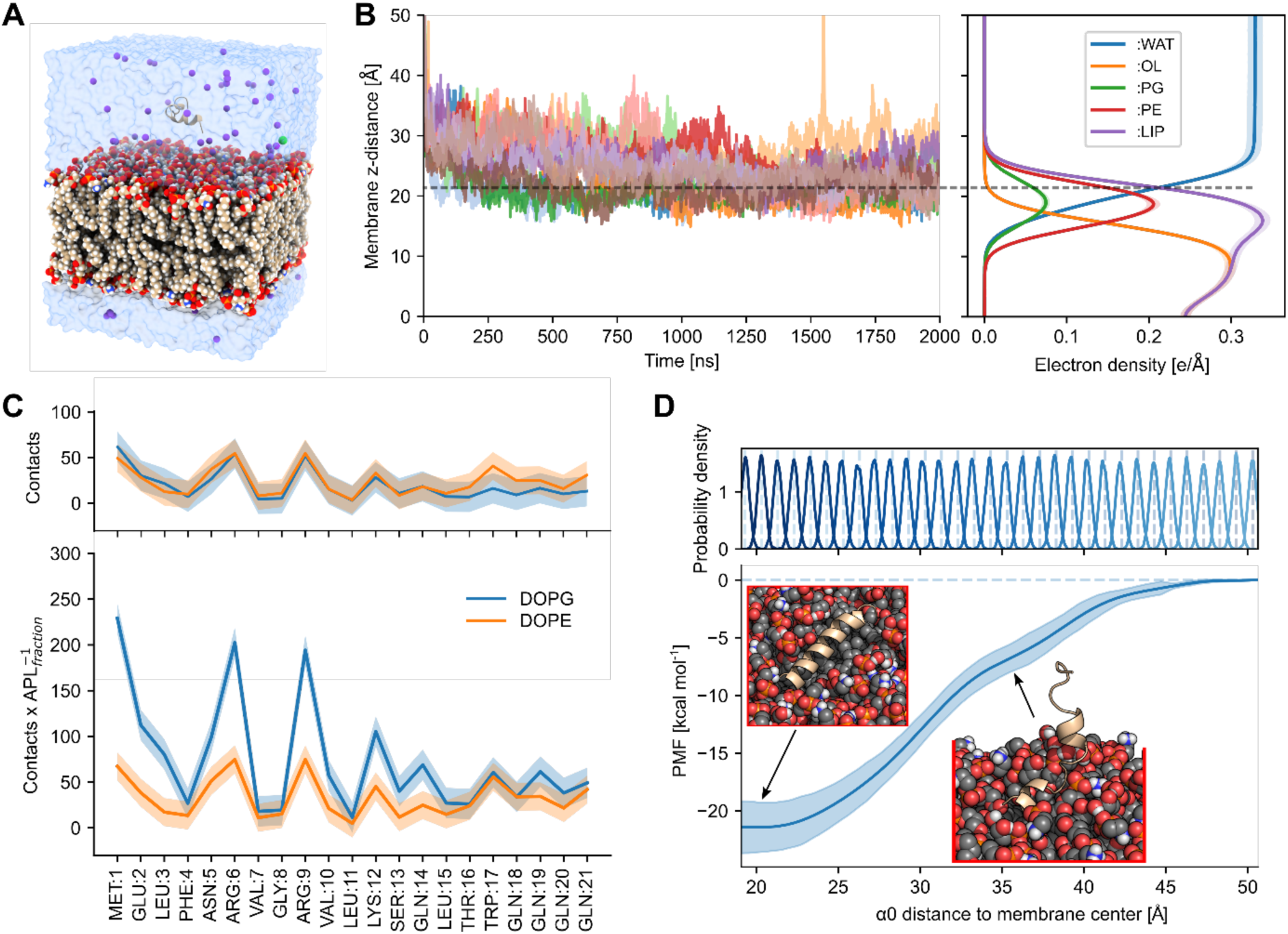
Binding of peptide α0 to a DOPE:DOPG lipid bilayer in molecular dynamics (MD) simulations. **A:** Starting configuration for MD simulations, helix *α*0 located ∼25 Å above a DOPE:DOPG (3:1) lipid bilayer, water, and K^+^ as counterions. **B:** *α*0 binds spontaneously to the membrane surface in less than 100 ns, as expected for amphipathic peptides. Left: distance of the center of mass of the peptide along the membrane normal. Right: electron density profile, showing a monolayer arrangement with water, headgroup, and inner membrane regions. PG, phosphatidylglycerol; PE, phosphatidylethanolamine; OL oleic acid tail; WAT, water; LIP, sum of all lipid densities. The dashed line indicates the region of the water-membrane interface. **C**: The average number of contacts of peptide residues with the lipid head groups shows that M1, R6, R9, and K12, all positively charged, interact more frequently with lipid headgroups than the other amino acids (top). Scaling the number of contacts with the area per lipid for DOPG and DOPE to consider the membrane composition reveals that interactions with negatively charged lipids (DOPG) are more frequent (bottom). **D**: From umbrella sampling simulations, using the distance to the membrane center along the normal as reaction coordinate, a free energy profile (PMF) for peptide (un)binding was computed. The shaded area shows the standard deviation using the unbound state as a reference (see Materials and Methods). The profile has a minimum at the membrane surface (∼20 Å, B right), with a depth of ∼-20 kcal mol^-1^, and increases with a slope change between 30 and 35 Å until a distance of 45 Å is reached. Insets show the last structures from umbrella sampling windows 2 and 17 (restrained at 20.3 and 36.3 respectively).

### Cryo-EM reveals dilated rods engulfing membranes through helix α0 interactions

To understand the mechanism of membrane remodeling by PspA, we examined cryo-EM structures of PspA in the presence of EPL membranes. For our analyses, we refolded PspA in the presence of 50 nm sized small unilamellar EPL vesicles, similar to a recently described protocol (Junglas et al., 2024a, 2024b). In the EPL sample, we detected a large number of PspA rods showing varying diameters within one rod, revealing the lack of a persistent structure along the rods (**Fig. 3A**). In the presence of EPL membranes, the rods have a wider diameter and a broader diameter distribution (200 to 345 Å) in comparison with the diameter values and distribution of the previously analyzed *apo* protein (180 to 290 Å) (**Fig. 3A**) (Junglas et al., 2024a). By segmenting the rods of the EPL-PspA sample and sorting the segments according to diameter through classification, we determined a total of 10 cryo-EM structures with resolutions ranging from 4.7 to 6.5 Å (**Suppl. Fig. 1A, B, Table 1**). While the imposed helical symmetry parameters suggest no architectural relationship between the PspA rods, helical lattice plots reveal that the left-handed helical rung corresponding to layer line ∼110 Å in the power spectrum increases in Bessel order from n=10, 11, 12…18 with increasing diameters of 200, 215, 235…345 Å, respectively (**Suppl. Fig. 1**, **Suppl. Table 2**). Upon insertion of an additional subunit into the helical rung, the assembly widens in discrete steps of around 20 Å up to a diameter of 345 Å. The rods with diameters larger than 280 Å were of particular interest as they showed enclosed double-layered lipid density in the cross-sectional top and side views in addition to the expected PspA protein densities, whereas rods of 270 Å diameter contained only partial lipid density and smaller diameters showed no density in the cross-section views. Due to the critical role of helix α0 in membrane interaction, we also prepared and refolded an N-terminally truncated PspA including helices *α*1-5 in the presence of EPL liposomes. For the truncated PspA *α*1-5 sample, the total diameter distribution was narrower and ranged from 235 to 290 Å, with two maxima at 255 Å and 275 Å that are closer but not identical to the *apo* distribution of WT PspA (**Fig. 3B**). In the PspA *α*1-5 + EPL sample, we determined a total of five cryo-EM structures with resolutions ranging from 3.8 to 5.4 Å supported by visible side-chain details (**Suppl. Fig. 1C, D**). None of these structures contained any additional lipid density in the rod lumen. Recent work has shown that PspA is an atypical ATPase with a low hydrolysis rate (Junglas et al., 2024a) while this ATPase activity has been suggested to relate to PspA’s membrane remodeling activity. Using an ADP-Glo assay, we confirmed the reported ATP hydrolysis by the WT protein with a rate of 3 h^-1^. Interestingly, when the protein was reconstituted in the presence of membranes, the ATPase activity was increased by ∼210 %, while the activity of PspA *α*1-5 was not affected in the presence of membranes (**Suppl. Fig. 1E**). The combined analyses of the isolated PspA helix α0 and the PspA cryo-EM structures suggest that for membrane engulfment into PspA rods helix α0 as well as the formation of rods with diameters beyond 280 Å are required and that PspA hydrolysis of ATP is boosted by the presence of membranes.

**Figure 3:**
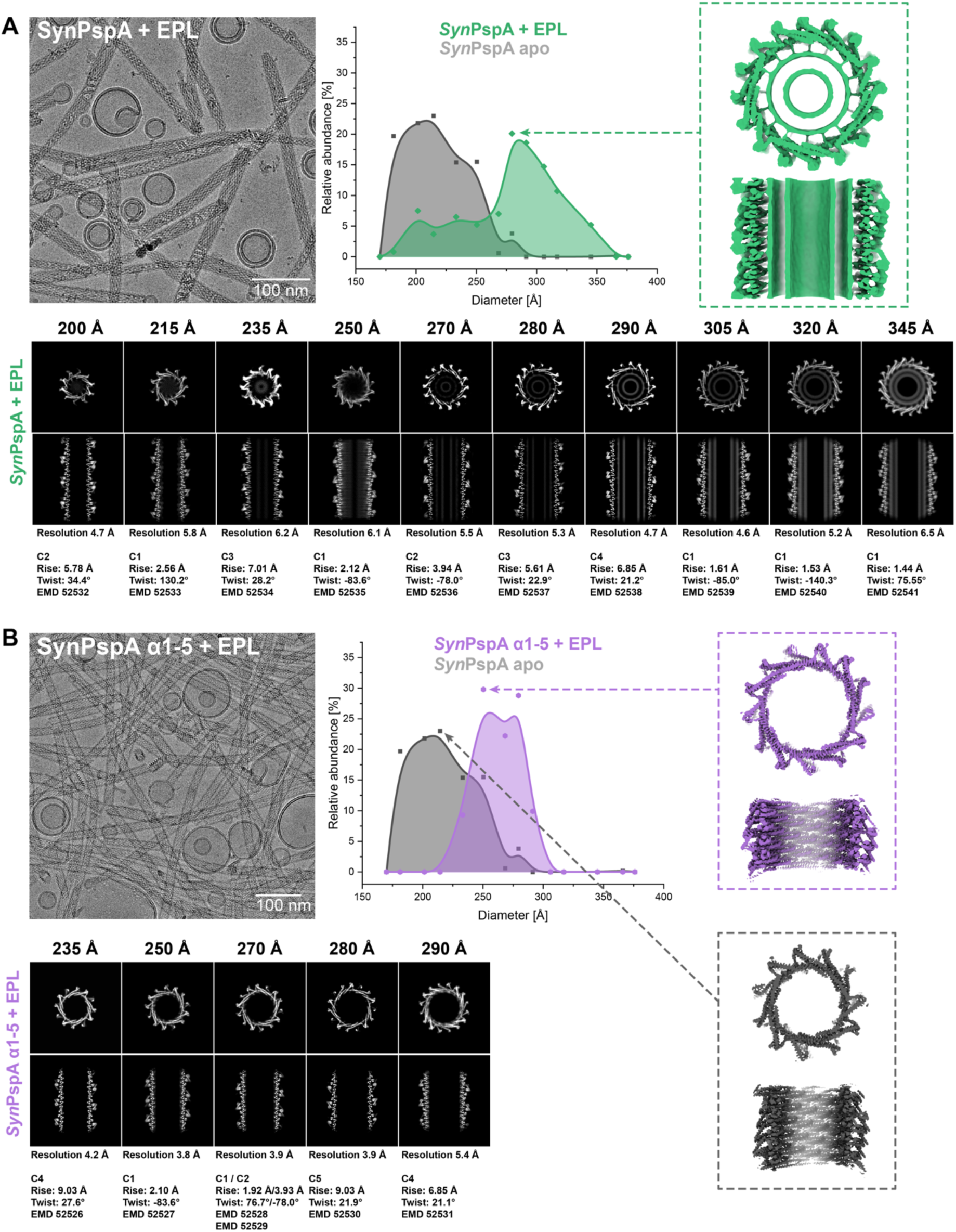
PspA rod diameter distribution and corresponding cryo-EM structures in the presence of membranes. Example micrographs (left) and PspA rod diameter distributions (center) for **A:** PspA+EPL (green), and **B:** PspA *α*1-5+EPL (violet), based on the relative occurrence of rod segments with a certain diameter. Highlighted structures (right) show details of the most abundant diameter for each sample (PspA+EPL (green), PspA *α*1-5+EPL (violet), PspA *apo* (grey)). A gallery of PspA rod cryo-EM structures (bottom) with cross-sectional top views or *z*-slices (top row) and cross-sectional side views or *xy*-slices (bottom row) is shown, with information on resolution and symmetry below.

To investigate the details of membrane interaction, we refined ten atomic models of the different rod diameter assemblies using the determined cryo-EM maps based on the previously published structure of PspA (PDB:7ABK). When comparing them with the previous *apo* structures (Junglas et al., 2024a, 2021), we observed that the structures of the EPL sample were identical to the *apo* structures with the same diameters, including their respective helical symmetry (**Fig. 4A** and **Supp. Fig. 5A**). For the *α*1-5 PspA variant, we found only five rod diameters (235, 250, 270, 280, 290 Å), whereas two different symmetries (C1 and C2) could be identified for the diameter of 270 Å (**Suppl. Fig. 1B**). In the WT structures, we were able to clearly assign EM density to helices *α*1-5 whereas continuous density corresponding the helix α0 could only be observed in diameters 280, 290, 305, 320 and 345 Å and not for the 270 Å diameter (**Fig. 4B**). Helix α0 was found as a well-defined density stalk that connected the core PspA-fold with the bilayer density, creating a constant distance of ∼35 Å between helix α0 and the bilayer center (**Fig. 4C**). This connecting stalk only accommodated half of the α0 helix before merging perpendicularly with the outer leaflet of the bilayer in the rod lumen. The density and the modeled helix α0 structure resembled the intermediately bound structure found in US simulations of isolated helix α0 (see **Fig. 2D**, inset). Therefore, we included the kink at G8 in helix α0, as identified in the US, which allowed the alignment of the positively charged residues R6, R9 and K12 with the negatively charged membrane headgroup region. Thus, the N-terminal part of helix α0 inserts into the lipid bilayer likely promoting positive membrane curvature. When comparing the tubulated EPL membranes inside rods, we found that the bilayer thickness remained constant (34 Å) regardless of the different rod diameters, even when the radius of the rods and thus the outer membrane leaflet radius increased (**Suppl. Fig. 1C-E**). Together, the PspA cryo-EM structures solved in the presence of membranes revealed structural details of how α0 binds to membranes and lipid interactions contribute to internalizing membrane tubules into the lumen of PspA rods with different diameters.

**Figure 4:**
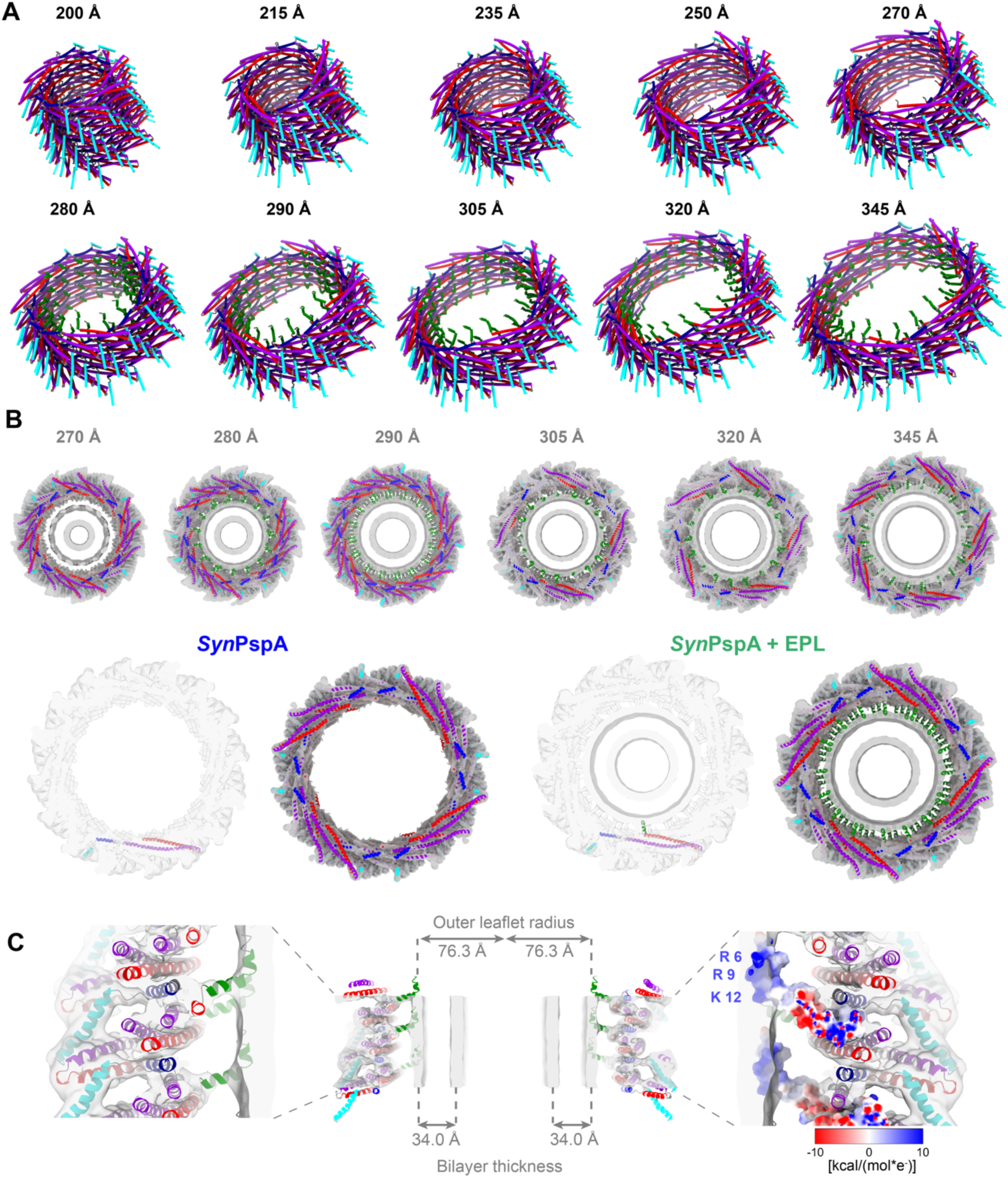
Membrane interaction of PspA rods. **A:** A total of 10 ribbon models of helical assemblies of PspA+EPL with diameters from 200 to 345 Å (helix α0 green, *α*1 red, *α*2+3 violet, *α*4 blue, *α*5 cyan). **B:** Top row: Gallery of PspA+EPL rods containing significant lipid density (top views of the cryo-EM density maps with the atomic model of the respective polymer structure, helix α0 green, *α*1 red, *α*2+3 violet, *α*4 blue, *α*5 cyan). Bottom row: Top view of the cryo-EM density maps with the atomic model of the 290 Å PspA rods from PspA (left) and PspA+EPL (right); helix α0 green, *α*1 red, *α*2+3 violet, *α*4 blue, *α*5 cyan. **C:** Sliced side view of a 290 Å PspA+EPL rod. Left: Vertical sliced side view of the cryo-EM density map with the atomic model. Center: Complete vertical density section with atomic model including measurements for the outer leaflet radius and the bilayer thickness of the engulfed lipid tube. Right: Sliced side view of the cryo-EM density map with the Coulomb electrostatic potential of the monomer. R6, R9, K12: Positively charged amino acids of helix α0 partially inserted in the headgroup region of the bilayer.

### Molecular dynamics simulations capture the contributions of helix α0 binding and membrane bending energy

The cryo-EM analysis revealed that only PspA tubes wider than 280 Å contained a clearly resolved membrane tube in their inner lumen (see **Fig. 3**). To investigate the structural dynamics and energetics of a representative 290 Å diameter PspA rod in the presence of a membrane, we performed coarse-grained (CG) simulations with the SIRAH force field (Barrera et al., 2019; Machado et al., 2019). Unbiased simulations over 10 µs of eight replicas showed no strong interaction of 160 Å high PspA assemblies with the membrane surface; in only three replicas the PspA assembly approached the membrane but did not bind to it (**Fig. 5A**). We thus resorted to biased CG simulations to steer the membrane through the center of the PspA rod. For this experiment, the replica that showed in its last frame the shortest distance along the membrane normal between the PspA assembly and the membrane was chosen. Two structures at different pulling points were backmapped to a full-atom representation: (*i*) where the membrane reached ‘half-way-through’ (HWT) the assembly and (*ii*) where the membrane reached ‘all-way-through’ (AWT), *i.e.*, fully covers the height of the PspA assembly (**Fig. 5B, Suppl. Fig. 1A**). An additional no-protein control’ based on full-atom AWT, was prepared, for which the protein was replaced by water to evaluate membrane behavior in the absence of PspA. In all eight PspA-containing unbiased MD simulations of 300 ns with a full-atom representation starting from AWT, the membrane tubule remained internalized in the lumen of the PspA rods. In contrast, when starting from HWT, the membrane retracted from the lumen in all eight replicas and became flat. The same retraction was observed for the control simulations of the AWT membrane shape when no PspA was present.

**Figure 5:**
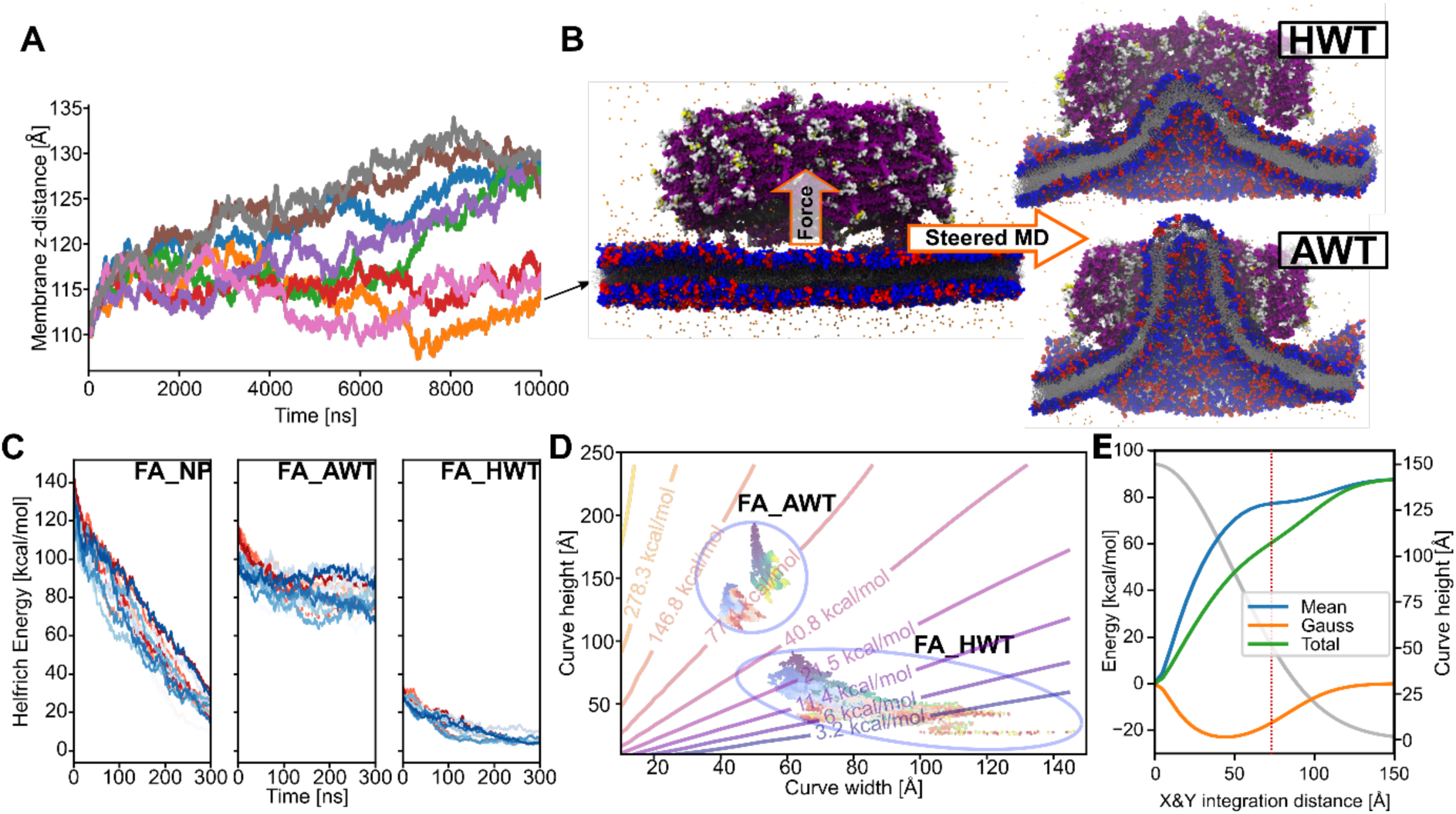
Interactions of the PspA 290 Å diameter assembly with the membrane. **A:** In coarse-grained molecular dynamics simulations (CG-MD), for three out of eight replicas the center of mass distance of PspA to the membrane center along the membrane normal decreased. **B:** The final structure with the shortest distance was used as a starting point for steered CG-MD. Two structures were selected to be backmapped to a full-atom representation and further analyzed, one with the membrane ‘half-way-through’ (HWT) and one with the membrane ‘all-way-through’ (AWT). **C**: Helfrich energy over the MD simulation time for full-atom no PspA (FA_NP), full-atom AWT (FA_AWT), and full-atom half-way-through (FA_HWT). The shape of the upper (shades of blue) and lower (shades of red) membrane leaflets in the simulations were approximated by 2D Gaussian functions, from where the curvature and the associated Helfrich energy were computed. (Also refer to **Suppl.** Fig. 6B). **D**: Helfrich energy isoenergy contour plot for membrane surfaces represented with different heights and widths as 2D Gaussian functions. The same width was taken for both dimensions. Helfrich energies of snapshots of FA_AWT and FA_HWT trajectories are projected onto the isocontours (blue to yellow, for start-to-finish upper leaflet, and light blue to red, for start-to-finish lower leaflet). For a 3D depiction of the energy surface refer to **Suppl.** Fig. 7. **E**: Total, mean and Gauss bending energy contributions as a function of integration limits in the X/Y plane from the center of the membrane surface. A representative Gaussian function for FA_AWT (see panel D) with 150 Å of amplitude and 50 Å of standard deviation in both dimensions was used (gray, right y-axis). The vertical dashed line shows the calculated inner cavity radius as calculated by HOLE (Smart et al., 1993) on the cryo-EM structure. The major energy contribution occurs close to the tip of the membrane surface, while increases become smaller as the inner diameter of the protein is reached.

Based on these calculations and observations, we hypothesized that two opposing energetic contributions give rise to the observed differential behavior of the PspA-membrane systems: on the one hand interactions between PspA and the membrane, mediated by helix α0 (see **Figs. 1 and 2**) that favor membrane internalization, and, on the other hand, energetic costs associated with bending a membrane that disfavors membrane internalization (Helfrich, 1973). Throughout the MD simulations starting from AWT, PspA tightly interacted with the curved membrane surface mainly through residues 1-81, with a major contribution from α0 and, in particular, R9 (**Suppl. Fig. 1B**). As the AWT simulations progressed, the number of α0 helices interacting with the membrane surface increased to 20-30 (out of 60) (**Suppl. Fig. 1C**) whereas in the case of HWT, less than ten α0 helices interacted with the membrane throughout the MD simulations. Assuming that the PspA-membrane interaction is dominated by each helix α0-membrane interaction, the favorable contribution to membrane internalization is thus at least two times larger in the AWT than in the HWT scenario. In support of this analysis, when we abolished the α0-membrane interaction by removing helix α0 lacking this binding energy contribution, we did not observe any membrane internalization in the lumen of the PspA (α1-α5) rods in the cryo-EM images (see **Fig. 3B**).

In order to assess the energy associated with bending a symmetric lipid bilayer, we used a Helfrich Hamiltonian that describes the energy in terms of the mean and Gauss curvatures at a given surface area segment (see Materials and Methods). To obtain a smooth representation of the membrane curvature caused by PspA adhesions, while removing local curvature effects due to fluctuations of the membrane surface, the shape of the leaflet surfaces was approximated by a 2D Gaussian (“bell curve”) function. The Helfrich bending energy computed for the ‘no-protein-control’ and HWT MD simulations approached zero over time (**Fig. 5C**), as expected from the decreasing curvature of the retracting membrane. In contrast, for the AWT simulation, the Helfrich energy decreased over the first half of the MD simulations but then plateaued after approximately 150 ns at values between 70 and 90 kcal mol^-1^ for the upper and lower leaflets, in line with the membrane tubule remaining internalized. Notably, in this case, the total Helfrich energy of ∼160 kcal mol^-1^ is within the range of the total binding free energy of 20-30 helices α0 to the membrane (∼125 to ∼190 kcal mol^-1^), suggesting that the relaxed AWT simulation was in an equilibrium state. To generalize this analysis, Helfrich energies were computed for membrane surfaces with different heights and widths represented as a 2D Gaussian function and displayed as isoenergetic lines (**Fig. 5D**). The Helfrich energy steeply rose when the curve height increased and when the curve width decreased (**Suppl. Fig. 1**). Helfrich energies obtained for snapshots of the AWT and HWT trajectories were projected onto the isoenergetic lines. In both cases, the scenarios started at higher Helfrich energies and relaxed over the simulation time towards membranes with lower heights and larger widths, although AWT isoenergetic lines remained in a region of medium Helfrich energy (see also **Fig. 5C**). Notably, the AWT line is also located in a region where the Helfrich energy rapidly increased with decreasing membrane widths. The energetic consideration suggests that PspA bending of membranes to narrower diameters is costlier than to wider diameters. The computed bending energies for narrower diameters may explain why PspA rods below a diameter of 280 Å were not found to engulf any membrane in the cryo-EM images: For rods with higher diameters the required bending energy is reduced, while in parallel the number of interacting α0 helices increases, which may explain the experimentally observed shift towards wider diameters (see **Fig. 3A**).

Moreover, integrating the membrane surface from the tip of the membrane surface to the rim in a stepwise manner and plotting it against the Helfrich energy reveals that the major Helfrich energy cost occurs next to the membrane-interacting tip indicated by a change in the steep slope (**Fig. 5E)** (see Materials and Methods). While the mean curvature component is highly unfavorable in this region, the Gauss curvature component contributes favorably, which has been shown to be a driving component in membrane fission of membrane tubules (see below) (Rueda-Contreras et al., 2021). Once the initial curvature is induced, the highest energy cost has been overcome, while extending the tubule is less costly and will be sustained by further binding of α0 helices in the lumen of the PspA rod. Overall, the higher bending energy cost for forming high membrane curvature suggests that wider initial diameters of PspA with less curved membranes require a lower cost for initiating membrane tubulation. MD simulations together with bending energy considerations of membrane tubule formation reveal the initially high bending energy costs for inducing high membrane curvature, while successive and additive binding of the N-terminal helix α0 to membranes interactions inside the assembly lumen finally overcome the required bending energy by binding energy gains, resulting in membrane tubulation inside tubular PspA assemblies.

### Visualization of membrane structures in variable diameter tubes

Next, we acquired tomograms of the PspA+EPL sample and comprehensively analyzed the PspA ultrastructures and associated membrane structures (**Fig. 6A)**. Membranes were found in the form of stand-alone vesicles, rod-attached vesicles and vesicles internalized in the lumen of PspA rods (**Suppl. Fig. 1A**). In agreement with the single-particle micrographs of PspA rods, a detailed analysis of the diameter distribution showed that most rods (with or without membrane) vary in diameter along their length, while only a few maintain a constant width (**Suppl. Fig. 1B**). When we counted vesicles attached to the rod ends, we found 106 (64 %) vesicles close to the thinner and 59 (35 %) vesicles close to the thicker end typically spanning diameters between 20 and 75 nm (**Fig. 6B**). When we quantified the number of rods containing tubulated membranes inside, 84 out of 249 (34 %) were located on the thick end of the rods, the remaining 165 (66 %) rods had vesicles centrally incorporated, whereas no vesicles were internalized at the thin end, in line with the above estimated lower Helfrich bending energy that needs to be overcome for wider rods (see **Fig. 5D**). Interestingly, only those vesicles attached to the thicker ends showed an internalized lipid bilayer as part of a continuous vesicular bilayer, in agreement with the cryo-EM structures that revealed internal lipid bilayer density for rods with diameters between 280 and 345 Å. Within the lumen of the rods, we identified in some cases continuous membrane tubules, while in other cases discontinuous membrane structures, i.e., PspA rods engulfed multiple tubulated vesicles as separated mini-vesicles, vesicular discs or less structured lipid assemblies (see **Fig. 6A**, red and green arrows, respectively). One or two tubular vesicles and two to five membrane discs occur in an average rod (**Fig. 6C** and **Suppl. Fig. 1C**). The segmented three-dimensional tomograms confirmed that these often densely packed membrane structures inside the PspA rods were well-separated and not part of a connected membrane network (**Fig. 6D**). When following PspA rods towards smaller diameters, the EPL membrane inside the lumen appears to be constricted concomitantly until separated membrane structures appear. In some cases, we found rods with membrane attached on both ends, first internalizing a continuous lipid bilayer resulting in membrane tubulation at the wider end of the rod, and a small vesicle attached to the thinner end as if it was just going to leave the rod (**Fig. 6E**). Interestingly, the emergence of separated membrane structures in the PspA rod lumen and next to the narrow end of membrane tubules is consistent with a previously discussed pearling mechanism of membrane fission (Rueda-Contreras et al., 2021).

**Figure 6:**
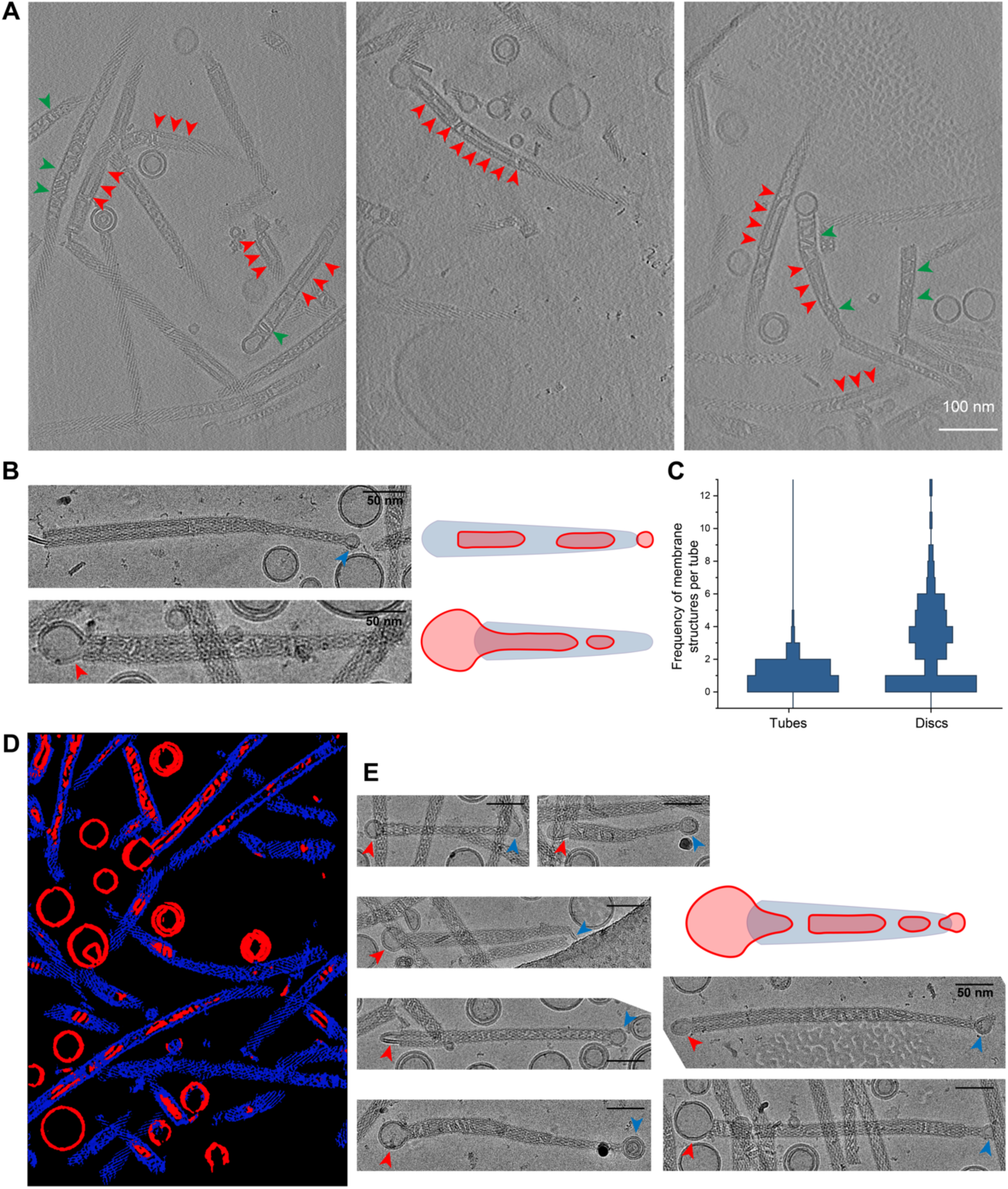
Tomographic analysis of PspA-remodelled membranes. **A**: Tomogram slices of PspA with EPL membranes. Red arrows: continuously tubulated EPL vesicles within the PspA rod structures. Green arrows: discontinuous membrane discs within the PspA rod structures. **B**: PspA frequently forms conically-shaped rods, with vesicles at the wide as well as the narrow end of the rods. Blue arrow: vesicles near the thinner end of the PspA rod structures do not reach into the lumen of PspA. Red arrow: vesicles near the wider end of the PspA rod structures are tubulated into the lumen of PspA. **C:** Bar plot of membrane tubes and membrane discs located within the PspA rods (n = 125) **D**: Segmentation of a PspA tomogram with tubulated EPL membranes. Blue: PspA rods. Red: EPL vesicles and tubulated membrane within thein PspA rod lumen. **E**: Red arrows: Beginning of membrane tubulation of EPL vesicles on the thicker side of the PspA rod structures. Blue arrows: vesicles near the thinner end of the PspA rod structures.

## Discussion

In this study, we investigated the interaction of the bacterial ESCRT-III protein PspA with membranes. We analyzed the role of helix α0 in membrane interaction by focused biophysical experiments and MD simulations (**Figs. 1, 2**). Using cryo-EM, we found the diameter distribution of PspA rod assemblies to be affected by lipid interactions, resulting in the formation of larger rod diameters that are capable of internalizing membranes (**Fig. 3**). The truncation of helix α0 reveals its critical role, as PspA α1-5 was no longer able to internalize and tubulate membranes. In the determined cryo-EM structures with internalized membranes present, we observed that α0 formed kinked α-helix when bound to negatively charged membranes (**Fig. 4**). MD simulations focusing on the interaction of PspA rod assemblies with a lipid bilayer revealed that overcoming the membrane bending energy presents a critical step for internalizing membranes into the lumen of PspA assemblies, indicating that wider rods energetically favor membrane initialization and tubulation, which explains why rods are found to be wider in the presence of membranes (**Fig. 5**). Continuous membrane tubules, thinned discontinuous mini-vesicles and lipid discs were found in the lumen of PspA assemblies often next to shedded vesicles at the thinner rod ends (**Fig. 6**).

In the bacterial ESCRT-III proteins PspA and Vipp1, the N-terminus can form a 20-30 aa α-helix (helix α0) in the presence of membranes (Junglas et al., 2021; McDonald et al., 2017; Osadnik et al., 2015). In eukaryotic ESCRT-IIIs, this extension tends to be shorter (3-20 aa) and has not been shown to form a structured α-helix to date albeit in the CHMP2A/CHMP3 structure, the corresponding residues are positioned toward the membrane interface (Azad et al., 2023). Yeast ESCRT-III proteins, *i.e.* Snf7, Vps24, Vps2, have amphipathic N-terminal extensions that can function as a membrane anchoring domain, mediating the contact of the ESCRT-III polymer to the membrane (Buchkovich et al., 2013). As shown for PspA and Vipp1, the hydrophobic face and positively charged residues are critical for membrane interaction of α0 (this study and (Gupta et al., 2021; Junglas et al., 2024b)). For bacterial ESCRT-III α0 helices, the observed conformation and membrane interaction mode appear to differ between PspA and Vipp1: While helix α0 in Vipp1 forms a straight helix that is embedded into the headgroup region of an outer membrane leaflet with its full length (Junglas et al., 2024b), helix α0 in the now solved PspA structures is kinked and only the first half of the helix is embedded in the headgroup region of the outer leaflet. Consequently, the distance between the outer membrane leaflet and rod wall also differs between PspA and Vipp1. In Vipp1, the outer leaflet is in contact with the inner ring/rod surface (*i.e.* α1 and 2/3), while in PspA, a gap of 35 Å between the membrane center and the rod wall does not indicate a direct interaction of α1+2/3 with the membrane. Here, helix α0 serves as a stalk between the protein and the membrane, maintaining a constant distance. These different α0 conformations likely also affect membrane binding of PspA and Vipp1, such that Vipp1 offers a much larger membrane interaction surface (full α0 and α1+2 in Vipp1 *vs.* only half of α0 in PspA). Furthermore, in Vipp1, helix α0 also interacts with the rod wall via its positively charged residues to stabilize certain assembly types (Junglas et al., 2024b). This is not the case in the PspA+EPL structure, where helix α0 does not show interactions with other parts of the protein. Interestingly, the Vipp1 structure showed an assembly type similar to PspA rods, when helix α0 was removed (Gupta et al., 2021; Junglas et al., 2024b; Thurotte and Schneider, 2019).

Furthermore, our ESCRT-III interaction data reveal strong similarities to membrane interactions of epsin and N-BAR proteins. Similarly, an amphipathic N-terminal helix (that is unfolded in the absence of lipids) is embedded into the membrane and serves as a membrane anchor, while the core of the protein serves as a scaffold for polymer assembly and membrane shaping. For example, the N-BAR protein endophilin has been shown to remodel membranes driven by membrane insertion of its amphipathic N-terminal helix, which also results in membrane tubulation (Bhatt et al., 2021; Cui et al., 2013). Epsin and other ENTH domain proteins are suggested to induce membrane curvature either by wedging of their N-terminal amphipathic helix into the membrane or by molecular crowding (Ford et al., 2002; Steinem and Meinecke, 2021). The interaction of N-BAR H0 with membranes has been studied extensively by CG and atomistic MD simulations (Cui et al., 2013), revealing hydrophobic protein moieties that interact with hydrophobic parts of the bilayer. Notably, membrane defects expose hydrophobic parts that are more common in curved membranes. In return, the binding of H0 to membrane defects recruits additional defects, making this process highly cooperative (Cui et al., 2011). A curvature sensing and generating function has also been suggested for the PspA and Vipp1 α0 helices (McDonald et al., 2015), thus it is tempting to speculate that PspA follows a similar mechanism for curvature generation as put forward for N-BAR H0. Apart from curvature sensing and curvature generation, CG simulations have also shown that N-BAR H0 is crucial for stabilizing the N-BAR lattice around the tubulated membrane by dimerizing with neighboring H0 helices, thus helping to form contiguous BAR domain strings (Cui et al., 2013). Although helix α0 is not directly interacting with neighboring α0s in our PspA rod structures, in Vipp1 tubes and rings, neighboring α0 helices directly interact with each other to form contiguous columns at the inside of the polymers that are suggested to stabilize certain assembly types (Junglas et al., 2024b).

In order to better understand membrane curvature induction and tubulation, it is particularly important to understand the energy costs necessary for inducing membrane curvature and how this may be provided through protein binding. In the case of PspA, one of the key events required for membrane tubulation likely is primarily mediated by the vertical insertion of the entire amphipathic helix α0 into the headgroup region of the outer bilayer leaflet. As observed in this manuscript by CD and Trp fluorescence spectroscopy and supported by conventional MD simulations, helix α0 binds to and forms an α-helix upon insertion into the membrane surface. The cryo-EM structures and MD simulations showed that only rods wider than 270 Å internalize and tubulate membranes. As indicated in the simulations, the energy required for membrane bending likely is too high for thinner rods to bend and internalize membrane tubules. Presumably, this is also the reason why we had to perform MD simulations starting with all-way-through rod-internalized membranes for the 290 Å-wide rods used here instead of starting with a rod bound to a flat membrane. The US simulations revealed that the interaction of helix α0 with membrane surfaces has a strong enthalpic component, but is counteracted by a reduction in entropy, resulting in an estimated association free energy Δ*G* = -6.32 kcal mol^-1^ per helix, a value close to the experimentally determined Δ*G*. As multiple α0 helices inside a PspA rod are involved in membrane binding, multivalency may lead to a superadditive effect as the decrease in translational and rotational entropy from the association of two bodies is reduced after initial binding, which should favor stronger interactions with the membrane surface. Importantly, our data revealed that the interaction of helix α0 with the membrane causes helix α0 to kink at the structure-breaking amino acid G8, which enables a targeted interaction of the positively charged residues (R6, R9, K12) with the negatively charged head groups of the membrane. Notably, in our simulation data helix α0 in the membrane-bound state is initially membrane-bound as a full α-helix, the kink structure observed in an intermediate state explains the kinked structure observed in the cryo-EM fittings.

The electron micrographs and 3D tomograms of the PspA structures solved in the presence of EPL membranes revealed interesting structural features in addition to the averaged PspA structures. The sample included vesicular spherical membrane structures devoid of protein while lipids were commonly found internalized into the lumen of the PspA rods in the form of membrane tubules. In addition to the continuous membrane tubules, discontinuous membrane structures were found inside the lumen of PspA rods. The occurrence of separated membrane structures within PspA rods is often accompanied by a thinning of PspA rod diameters until in some cases they appear to emerge from the thinner rod ends. The structures observed are consistent with the pearling of small vesicles from the tip of membrane tubules that may be a result of spontaneous membrane fission (Rueda-Contreras et al., 2021). Although the previous studies investigated theoretical models solely based on the shape and the associated energies of the membranes in the absence of proteins, the similarities to our experimental observations in the presence of PspA are noteworthy. Likely, PspA rods provide the environment to cast membranes into the required shapes until they undergo vesicle shedding and subsequently leave the rods.

To summarize our observations, we propose the following model for PspA-induced membrane tubulation and vesicle fission: The process of tubulation is initiated by assembling complexes wider than 290 Å (or with small subunits that assemble at the membrane to form a complex wider than 290 Å) on membrane surfaces to minimize the energy barrier required for the induction of membrane curvature. Due to the positively charged residues of the N-terminal region, PspA readily binds to the membrane surface with the entire helix α0. This binding facilitates local curving of the membrane at the tip of an emerging membrane bulge. The energy released by the additive interactions of multiple helices α0 with the membrane compensates for the energy required to initially bend the membrane (**Fig. 7**, I). Furthermore, the high energy required for initiating membrane bending might also be provided by the energy released due to PspA oligomerization. Oligomerized PspA then stabilizes the membrane curvature; as more α0 helices interact with the membrane, more energy is gained that generates a pulling force leading to the formation of an emerging membrane tubule in the inside of the PspA assembly (**Fig. 7**, II). Due to the spatially restrained interactions in the lumen of the assembly with the tubulated membrane, helix α0 is partially pulled away from the membrane, which causes a kink in the helix around G8 that contributes to maintaining the membrane interaction of R6, R9, and K12 (**Fig.7**, III). Helix α0-membrane interactions in addition to the PspA homo-oligomerization lead to the growth of the tubules, once the bending energy has been overcome during tubule initiation. The distal ends of rods that contain internalized tubulated membrane, i.e. the tips of growing PspA rods, tend to thin towards smaller diameters. Towards the smaller diameters, vesicular membrane structures appear to spontaneously fission from a growing tubule tip, in analogy to the described pearling, until they can leave the rods at the thinner end (**Fig.7**, IV) (or they can be ejected at the proximal end, which leads to ILV formation by inward-vesicle budding (see (Junglas et al., 2024a, 2021)). The described vesicle-shedding pathway through the PspA rod is reminiscent of an ejection process through a molecular nozzle.

**Figure 7:**
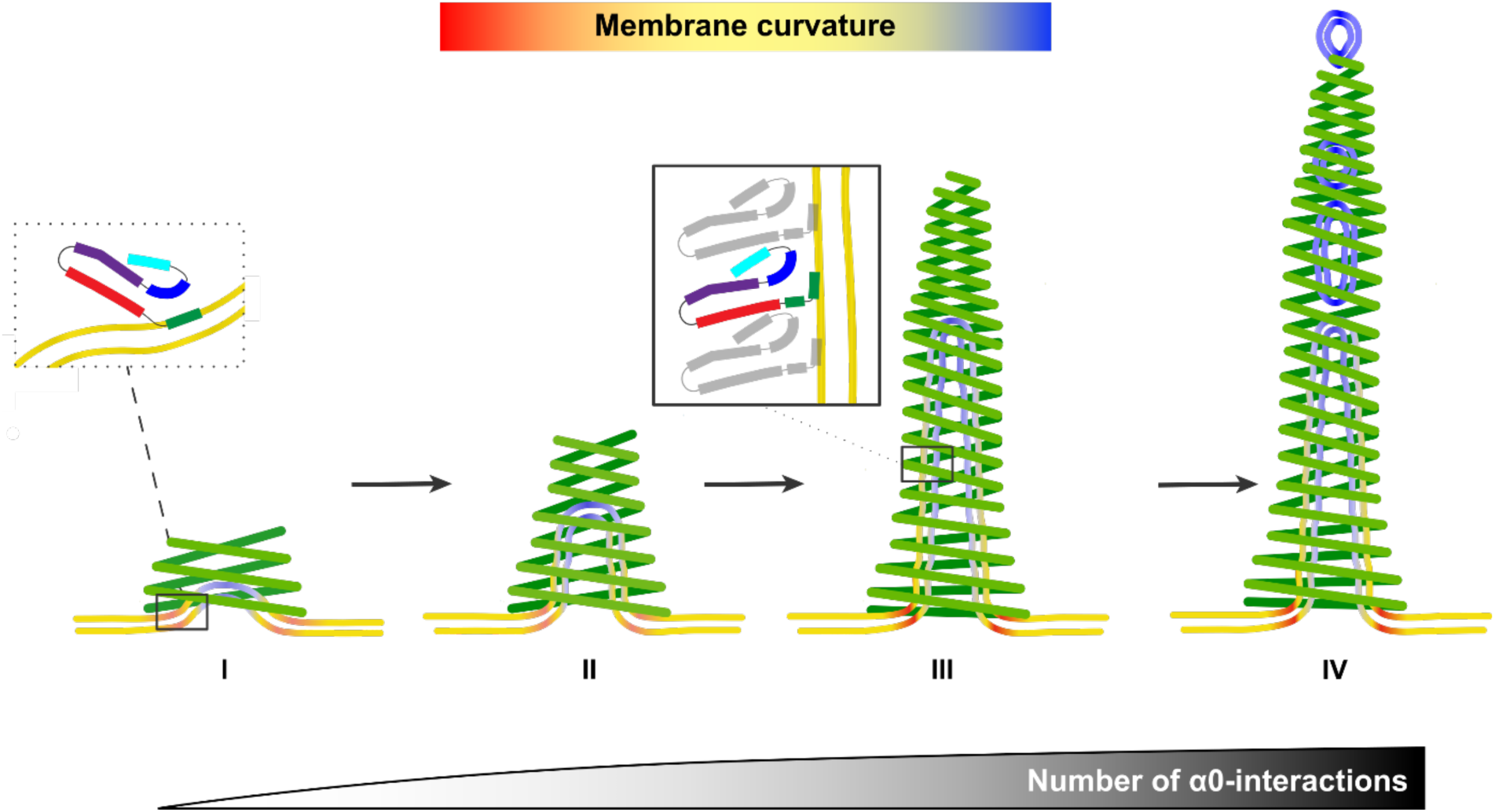
Model of membrane tubulation and vesicle fission mediated by PspA rod structures. **I**: PspA accumulates on the membrane surface and the complete helix α0 inserts into the membrane surface (inset). Complexes with a diameter > 290 Å are formed and initiate a low-curvature tubulation. **II**: The initial interactions allow further recruitment of PspA subunits into the complex leading to additional helix α0 interactions with the membrane, which propagate an emerging membrane tubule in the rod lumen. **III**: Upon extension, the membrane becomes fully tubulated in the lumen of a PspA rod. The membrane has the highest curvature at the tubule tip and base. Extension of the PspA rod with internalized membrane tubes involves partial unbinding of helix α0 by kinking at the structure-breaking amino acid G8 (inset), and as a result, solely the N-terminal part of helix α0 remains membrane-bound. Helix α0 green, α1 red, α2+3 violet, α4 blue, α5 cyan. **IV** As observed in tomograms (Fig. 6), in narrow ends of the PspA complex, smaller lipid vesicles emerge from the tubules upon fission of the membrane bilayer. The coloring of the bilayer schematically shows the mean curvature of the surfaces upon tubulation and fission; yellow indicates zero mean curvature.

While it may be tempting to compare the observed membrane thinning and vesicle shedding to other ATP/GTP-mediated membrane constriction machines such as dynamin (Hinshaw and Schmid, 1995), we emphasize that no ATP/GTP was required for the observed membrane remodeling albeit we had previously found that ATP can enhance the efficiency of PspA membrane deformation (Junglas et al., 2024a). The membrane thinning and vesicle shedding in the absence of ATP described here is merely a result of the energy gains through helix α0 binding, the directional internalization of lipid tubules towards smaller diameters, and the subsequent increase in negative Gauss curvature resulting in the spontaneous physical separation of small vesicles. The formation of these small vesicles can be explained in terms of a change in curvature energy driven by the Gauss curvature. The increase in membrane curvature favors fission events, as the transition from a single continuous membrane to vesicles is energetically favored by a negative Gauss curvature component (see **Figure 5E**) (Kozlov et al., 2010). Eventually, this term leads thin membrane tubules to spontaneously pearl in the absence of additional components (Rueda-Contreras et al., 2021).

Given the observed variation in PspA rod diameter with respect to membrane deformation, an important question arises regarding the associated dynamics: Do the rods act as a fixed-scaffold assembly or do they taper and widen dynamically? As the static cryo-EM images were plunge-frozen in time, we cannot conclude from the data presented here alone whether the PspA rod diameters maintain a fixed scaffold after tube assembly or whether they taper and widen dynamically. Previous studies revealed that the diameter distribution of PspA rods and the membrane binding capabilities were much smaller when comparing preformed rods with “in-situ” polymerized rods after refolding as used here (Junglas et al., 2024a, 2021). Thus, we can speculate that the rod diameter is predominantly defined during polymer assembly, and the membrane deformation observed here was rather a result of scaffolding and not dynamic tapering and widening. Further experiments with temporal resolution are needed to record rod diameters over time to clarify this question. An additional consideration is that the (re)assembly process may be highly dynamic in the cell when interaction partners such as chaperones or PspF constantly disassemble/assemble PspA rods, effectively leading to a dynamic tapering/widening process as suggested previously (Junglas et al., 2021).

We assume that PspA is involved in an active repair mechanism in the bacterial host cell. Our data now indicate that either PspA rods engulf damaged inner membrane patches by forcing the membrane into high positive curvature eventually leading to extraction of the damaged membrane and membrane re-sealing, or damaged membranes might be repaired by receiving membrane lipids through PspA-mediated vesicle shedding. How the proposed processes for bacterial ESCRT-IIIs compare with eukaryotic ESCRT-III activities, where vesicles are budded away from the cytosol in the case of canonical ILV formation, remains to be established. Several proposed models of eukaryotic ESCRT-III activities favor the stabilization of negative membrane curvature eventually, in collaboration with Vps4, membrane cleavage (Maity et al., 2019) directed away from the cytosol, resulting for instance in budding ILVs away in multivesicular bodies (Henne et al., 2011; Pfitzner et al., 2020). Although the exact mechanistic details remain unclear, several reports provide evidence that ESCRT-III filaments assemble within membrane tubes either *in vitro* (Azad et al., 2023; Bodon et al., 2011) or *in vivo* (Cashikar et al., 2014; Hanson et al., 2008). In contrast, PspA, Vipp1, archaeal CHMP4-7 and eukaryotic CHMP1B/IST1 have been shown to induce positive membrane curvature (Junglas et al., 2021; Liu et al., 2021; McCullough et al., 2015; Melnikov et al., 2022). Nevertheless, eukaryotic ESCRT-III are thought to be more versatile as they constitute heteropolymers formed by multiple different ESCRT-III subunits that interact in a certain sequence. So far, it has been shown that these heteropolymers bind to membranes and can tubulate and deform them due to the formation of such complex structures (Moser von Filseck et al., 2020; Pfitzner et al., 2020).

## Material and Methods

### Expression and purification of PspA

PspA WT (*orf slr1188*) of *Synechocystis* sp. PCC 6803 and associated mutants (*α*1-5 (deletion of *α*0 (aa 1-23)) were heterologously expressed in *E. coli* C41 cells in TB medium using a pET50(b) derived plasmid. For purification of PspA and associated mutants under denaturing conditions, cells were resuspended in lysis buffer containing 6 M urea (10 mM Tris-HCl pH 8.0, 300 mM NaCl) supplemented with a protease inhibitor. Cells were lysed in a cell disruptor (TS Constant Cell disruption systems 1.1 KW; Constant Systems). The crude lysate was supplemented with 0.1% (v/v) Triton X-100 and incubated for 30 min at RT. Subsequently, the lysate was cleared by centrifugation for 15 min at 40,000 g. The supernatant was applied on Ni-NTA agarose beads. The Ni-NTA matrix was washed with lysis buffer and lysis buffer with additional 10 – 20 mM imidazole. The protein was eluted from the Ni-NTA with elution buffer (10 mM Tris-HCl pH 8.0, 1000 mM imidazole, 6 M urea). The fractions containing protein were pooled, concentrated (Amicon Ultra-15 centrifugal filter 10 kDa MWCO), and dialyzed overnight against 10 mM Tris-HCl pH 8.0 (8 °C, 10 kDa MWCO) including three buffer exchanges. Protein concentrations were determined by measuring the absorbance at 280 nm of PspA diluted in 4 M guanidine hydrochloride using the respective molar extinction coefficient calculated by ProtParam (Gasteiger et al., 2005).

### Liposome preparation and membrane reconstitution

Chloroform dissolved *E. coli* polar lipid (EPL) extract was purchased from Avanti polar lipids. Lipid films were produced by evaporating the solvent under a gentle stream of nitrogen and vacuum desiccation overnight. The lipid films were rehydrated in 10 mM Tris-HCl pH 8.0 by shaking for 30 min at 37 °C. The resulting liposome solution was subjected to five freeze-thaw cycles, combined with sonication at 37 °C in a bath sonicator. SUVs (small unilamellar vesicles) were generated by extrusion of the liposome solution through a porous polycarbonate filter (50 nm pores). For PspA membrane reconstitution, unfolded PspA (in 6 M Urea) was added to EPL SUVs and incubated at RT for 15 min. Then the mixture was dialyzed overnight against 10 mM Tris-HCl pH 8.0 (8 °C, 10 kDa MWCO) including three buffer exchanges.

### Membrane binding of the peptides analyzed by CD and Trp fluorescence spectroscopy

The peptides PspA *α*0 WT [MELFNRVGRVLKSQLTHWQQQQEA] and PspA *α*0 RRAA [MELFNAVGAVLKSQLTHWQQQQEA] were purchased from PSL GmbH (Heidelberg, GER). This sequence corresponds to the first 24 amino acids of the *Synechocystis* PspA protein. Membrane binding of the peptides was investigated via monitoring Trp fluorescence changes. 2 µM peptide (10 mM Tris, pH 8.0) was incubated with increasing amounts of liposomes up to 200 µM lipid. The liposomes were prepared as described above. Upon incubation at 25°C for 15 min, corrected Trp fluorescence emission spectra were recorded using a fluorescence spectrometer (FP-8500, Jasco, Pfungstadt, Germany), from 300 nm to 450 nm with an excitation wavelength of 280 nm and an increment of 0.2 nm. The excitation and the emission slit widths were set to 2.5 nm. Buffer spectra were subtracted. The influence of lipid binding on the secondary structure of the peptides was determined via circular dichroism (CD) spectroscopy using a J-1500 CD-spectrometer (Jasco, Pfungstadt, Germany). 34 µM peptide (10 mM Tris, pH 8.0) was incubated with increasing amounts of liposomes, with lipid concentrations set to yield a similar lipid/peptide ratio as in the fluorescence spectroscopy measurements. After incubation at room temperature for 15 min, spectra were recorded in 1 nm steps at 25°C. Three individual samples were measured for each sample; 12 spectra were averaged. The secondary structure content of the peptides was calculated based on the CD spectroscopy data using BeStSel (Micsonai et al., 2022).

Peptide/membrane binding was quantitatively analyzed by fitting a hyperbolic binding model to the data, expressing the fraction of bound peptide as a function of lipid/peptide ratio r:

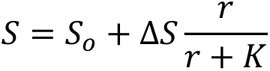

with *S* either reflecting fluorescence intensity or ellipticity. Thus, *K* is related to the respective *K*_D_ value via *K*_D_ = *K c*_o_, with *c*_o_ reflecting the peptide concentration in the given experiment (2 µM and 34 µM, respectively). The binding free energy calculations are based on the *K*_D_ values, with Λ*G*°=*RT*\*ln(*K*_D_), with *K*_D_ in M^-1^.

### All-atom conventional MD simulations of helix α0 peptide

The first 21 residues of the α0 peptide were taken from the first chain of the 290 Å diameter rod structure and capped in the C-terminus with an *N*-methyl group. The latter is required to avoid charge repulsion. The peptide was packed into a water/membrane-bilayer box using PACKMOL-Memgen (Martínez et al., 2009; Nugent and Jones, 2013; Schott-Verdugo and Gohlke, 2019) as included in AmberTools 23 (Case et al., 2023), using a membrane composition of DOPE:DOPG 3:1 and a minimum distance to the box boundaries of 25 Å, placing the peptide 25 Å above the membrane surface, as estimated by MEMEMBED (Nugent and Jones, 2013). The packed structure resulted in over 75,000 atoms in a box with dimensions 88.9 Åx88.9 Åx112.8 Å. The system was parameterized using the force fields ff14SB (Maier et al., 2015) for the peptide and LIPID21 (Dickson et al., 2022) for the lipids, and the TIP3P water model (Jorgensen et al., 1983). Hydrogen Mass Repartitioning (Hopkins et al., 2015) was applied, allowing to use a timestep of 4 fs.

Twelve replicas were stepwise relaxed, alternating steepest descent/conjugate gradient energy minimizations with a maximum of 20,000 steps each, using the pmemd.MPI implementation. The positions of the membrane were initially restrained during minimization; the final round of minimization was performed without restraints. During the relaxation process, all covalent bonds to hydrogens were constrained with the SHAKE algorithm (Ryckaert et al., 1976) within the pmemd GPU implementation (Le Grand et al., 2013). A direct space non-bonded cutoff of 10 Å was used. The Langevin thermostat with a friction coefficient of 1 ps^-1^ was used, while the pressure, when required, was maintained using a semi-isotropic Berendsen barostat (Berendsen et al., 1984) with a relaxation time of 1 ps, coupling the membrane (xy) plane. The system was heated by gradually increasing the temperature from 10 to 100 K for 5 ps under NVT conditions, and from 100 to 300 K for 115 ps under NPT conditions at 1 bar. The thermalization process was continued until 5 ns under NPT conditions were reached, after which production runs of 2 μs length were performed, using the same conditions. Trajectory coordinates were recorded every 200 ps in all cases.

### Adaptive steered and umbrella sampling MD simulations

In all replicas, the peptide bound quickly to the membrane surface (see main text). From the last frame of the first replica, adaptive steered molecular dynamics (AsMD) simulations were started, using the distance along the z-axis between the center of mass (COM) of the C_α_ atoms of the peptide and the COM of the phosphorous atom of the phospholipid bilayer as the reaction coordinate (RC) with a force constant of 100 kcal mol^-1^ Å^2^. Twelve iterations of pulling the peptide were performed, increasing the RC distance by 2.5 Å over 10 ns in each (0.25 Å ns^-1^), from 20.3 to 50.3 Å. In each iteration, 40 replicas were performed in parallel, selecting the one with a work profile closest to the Jarzynski average, as previously described (Ahmad et al., 2021; Bureau et al., 2015) his process allows to obtain a trajectory of a low free energy from the bound to the unbound state. To further obtain a free energy profile by means of umbrella sampling (US), intermediate structures were extracted every 1 Å and another window closer to the membrane center was added, resulting in 32 windows at a z-axis distance between 19.3 and 50.3 Å. The peptide was restrained with a harmonic potential at the corresponding RC value of a window using a force constant of 5 kcal mol^-1^ Å^2^. 1 µs of US was used per window, recording the RC every 4 ps. Samples showed a low correlation, as the highest autocorrelation time was 40 ps, as tested for window 3. The free energy profile or potential of mean force (PMF) was calculated by using the weighted histogram analysis method as implemented in the WHAM program version 2.0.11 (Grossfield, 2024). The final PMF and its error were obtained by taking the samples of the last 500 ns, averaging and calculating the standard deviation by generating 50 independent PMF calculations with 10 ns chunks and setting the value at the water bulk to zero.

### Free energy estimation from potential of mean force

For calculating the free energy of binding, the adsorption coefficient *κ* to the membrane surface was described by a modified approximation taken from Ben-Tal et al., 2000.

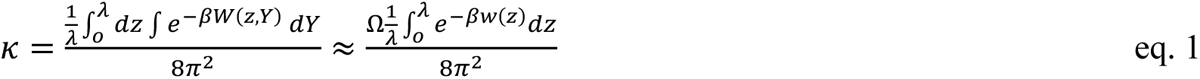

where *W* is the binding potential, which depends on the distance to the membrane (*z*) and the orientation of the peptide as described by Euler angles (*Y*, with pitch, roll and yaw being *θ*, *φ* and *ψ*, respectively) and λ is the membrane distance that defines a bound state. To calculate the pitch and roll angles, two reference vectors were constructed: a vector from the peptide’s center of mass to the center of mass of the first residue (v_1_) and one from the center of mass between the C_α_ atoms of residues 2 to 5 to the C_α_ atom of residue 2 (v_2_). The pitch was calculated as the angle between v_1_ and the membrane normal, while the roll was calculated by aligning v_1_ onto x^ and calculating atan2(v_2ž_,v_2ŷ_), where v_2ž_, v_2ŷ_ are x and y components of v_2_. By binding to the membrane surface, three of the six global degrees of freedom of the peptide are constrained, the position along the membrane normal and the pitch and the roll with respect to the membrane normal (**Suppl. Fig. 1C and D**). The calculated PMF *w*(*z*) captures the free energy as a function of the distance to the membrane, but not the dependence on *Y*. To consider the constraint in the pitch and roll, the bound state range was considered by *Ω* on the right side of eq. 1

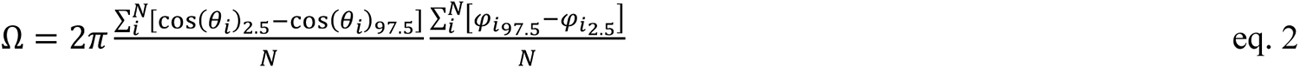

where 2π represents that any yaw angle is possible, the center term is the average span of pitch angles and the right term is the average span of the roll angles sampled on the windows in the bound state with *N* states, with the values 2.5 and 97.5 depicting the corresponding percentile values. Similar approaches have been used to estimate the impact of binding on rotational entropy (Ben-Shalom et al., 2017). As almost 100% of the integral of 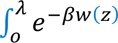 *dZ* is recovered when using windows that consider a RC ≤ 25 Å (**Suppl. Fig. 1G**), the first eight windows with RC values restraint between 19.3 and 27.3 Å were used, resulting in *Ω* = 2.19.

From *κ*, the binding free energy is obtained (Ben-Tal et al., 2000)

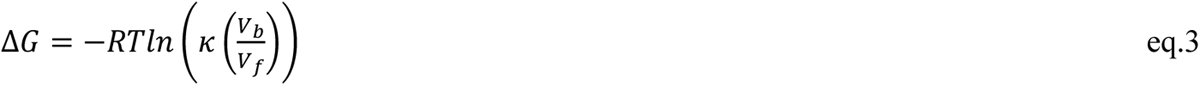

where *V*_b_ and *V*_f_ correspond to the bound and free volumes, respectively. Considering the standard state volume *V*_f_ = *V*_o_ = 1661 Å^3^, which corresponds to a concentration of 1 M, that the interaction with the membrane only constrains one translational dimension (Finkelstein and Janin, 1989) and that the bound distance along the membrane normal is λ, *i.e.*, *V*_b_ = *λ A*, with *A* being the membrane surface area,

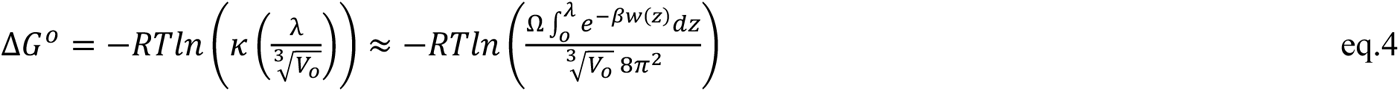

Eq. 4 results in Δ*G*^0^ = -18.55±2.07 kcal mol^-1^ for *T* = 300 K, which is in a similar order of magnitude as the distance term of a recent thorough multiple collective variable calculation for Phospholipase C, a peripheral membrane protein, but not as the overall binding free energy (Moutoussamy et al., 2022). This is because the approximation reflects the binding of a rigid peptide, considered a rigid cylinder, to the membrane surface (as explained in (Ben-Shaul et al., 1996)), but does not take into account the contribution due to conformational changes that happen to the peptide upon binding. In this case, the amphipathicity of helix α0 leads to a helix formation upon binding, which is entropically disfavorable, as the number of available conformational states is reduced. Furthermore, this effect is not captured by the distance reaction coordinate. At the plateau in the PMF, the sudden increase in reachable conformations does not cause a decrease in the free energy, as it becomes independent of the distance to the membrane surface (**Fig. 2D and Suppl. Fig. 1B-D)**. To estimate this contribution, the *k* nearest neighbors approach of the method PDB2ENTROPY (Fogolari et al., 2018) was used to estimate the entropy of residue torsions. For the analysis, we considered as bound structures windows 1 to 25, i.e., with restrained distances to the membrane center between 19.3 and 43.3 Å, and unbound structures those of windows 26 to 32, i.e, restrained between 44.3 and 50.3 Å. The rationale for this selection was that structures in windows 26 to 32 represent those mostly at the plateau of the PMF profile (**Fig. 2D**) and show a clear detachment along the trajectories, as evidenced by the increase in RMSD with respect to the bound structure and change in Euler angles (**Suppl. Fig. 1B-D**). This results in *T*Δ*S*_unbinding,conf_. = 12.23 ± 0.04 kcal mol^-1^ for *T* = 300 K (**Suppl. Fig. 1F**). Together with the result of eq. 4, the standard binding free energy of helix α0 is estimated as Δ*G*^0^_binding_ = -6.32±2.07 kcal mol^-1^.

### SIRAH coarse-grain simulations of a 290 Å diameter PspA complex

As for the helix α0 system, the PspA complex, consisting of 60 protomers, was packed using PACKMOL-Memgen (Schott-Verdugo and Gohlke, 2019), applying options to coarse-grain and parametrize the system with SIRAH (Barrera et al., 2019; Machado et al., 2019). The protein was protonated using PDB2PQR (Dolinsky et al., 2004) with predictions from PROPKA3 (Olsson et al., 2011). The structure was oriented using PPM3 with a flat Gram-negative membrane model (Lomize et al., 2022). As the phosphatidylglycerol (PG) head group is not available in the SIRAH force field, phosphatidylserine (PS) was used instead, which also has a -1 net charge, using a membrane composition of DOPE:DOPS 3:1. At the CG level, we do not expect major differences between both head groups. Considering that the membrane can be curved inside of the protein structure, extra padding was added to the system, ensuring at least 50 Å from the protein to the box limits, resulting in a system with dimensions of 426.6 Å x426.6 Å x201.8 Å and over 700,000 CG particles. Eight replicas of the system were prepared and minimized and relaxed in the same way as for helix α0, but totaling 50 ns in the thermalization step. As in the SIRAH AMBER tutorials, the following parameters were used: 20 fs timestep, Langevin thermostat with a collision frequency of 5 ps^-1^ set at 310 K, a semiisotropic Berendsen barostat (Berendsen et al., 1984) coupling the xy-plane with a relaxation time of 8 ps and a 12 Å direct space cutoff, using otherwise identical options as for helix α0. Production runs yielded 10 µs per replica.

### Membrane pulling through 290 Å complex

To obtain structures where the membrane interacts with the center of the PspA complex, the CG replica that showed the lowest distance between the center of mass of the complex and the membrane center after the production runs was selected. Similar to the pulling of helix α0 (see above), AsMD simulations were started, using as reaction coordinate (RC) the distance along the z-axis between the COM of the GC beads (C_α_ atom-equivalent) of the protein complex and the COM of the lipids that have any atom within 290/2 Å in the xy plane to the GC COM. Twenty iterations of 25 ns with 8 replicas in parallel were performed, selecting as a restart structure after each iteration the replica with the work closest to the Jarzynski average, pulling from a RC of -117.9 to 7.1 Å (0.25 Å ns^-1^). This results in a trajectory where the membrane fully reaches through the center of the protein complex.

### Backmapping of CG 290 Å diameter PspA / membrane structures and all-atom MD simulations

Two intermediate structures of the 290 Å complex with the membrane pulled were selected for all-atom simulations, one with the membrane ‘half-way-through’ and another with the membrane ‘all-way-through’ (HWT and AWT, see main text). To simulate at the all-atom level, the coordinates have to be properly backmapped from the CG level. For the protein, the VMD SIRAH tools plugin was used (Machado and Pantano, 2016). For the lipids, on the other hand, no automated solution is available for SIRAH lipids, partially, because the positions of the full-atom representation are ill-defined by the CG beads. To accomplish the backmapping, the following algorithm was implemented:

1. Coordinates of each lipid were extracted independently as PDB files.
2. Each bead was renamed according to the AMBER Lipid21 equivalent, as defined in the amber_lipid.map file included in AmberTools 23. PS residues were backmapped to PG residues, to recover a proper Gram-negative membrane.
3. Each lipid was parametrized with LEaP, letting the program add missing atoms according to the force field definition.
4. Each lipid was minimized independently with pmemd for a maximum of 5000 steps, with 2500 steps of steepest descent minimization and using igb = 1. All coordinates that were directly backmapped from the CG structure were restrained with a positional restraint of 5 kcal mol^-1^ Å^2^. To ensure proper structural isomers and proper stereochemistry, additional NMR restraints were imposed on acyl chain double bonds and the glycerol moiety of the head group.
5. The backmapped protein, all minimized lipids and ions were concatenated to reconstruct a backmapped system only lacking water.
6. The system was reparametrized with LEaP, imposing the box dimensions from the original CG simulation box.
7. A minimization of the concatenated system was performed with pmemd.MPI, imposing positional restraints on the previously restrained lipid atoms, ions and N/O/C_α_ backbone atoms of the protein, for 2500 steps of steepest descent and conjugated gradient minimization, respectively.
8. The minimized structure was resolvated using Packmol (Martínez et al., 2009), adding 11 water molecules for each WT4 residue in the original CG system, as expected for the SIRAH water model (Darré et al., 2010). A box smaller by 0.5 Å in each dimension was used, to avoid potential overlaps with periodic images.

The resulting structures were reparametrized with LEaP and simulated using the same simulation parameters as for helix α0, totaling over 5,380,000 atoms. Due to the system size, the minimization protocol used for helix α0 was not practical. At the same time, as the protein and membrane structures come mostly from stable CG simulations and previous minimization runs, mostly water clashes had to be removed. For this, a single pmemd.cuda minimization run with SHAKE enabled was performed, allowing a maximum of 10,000 steepest descent/conjugated gradient steps. Further thermalization and simulation conditions were the same as those used for helix α0 (see above). Eight production replicas of 300 ns length were performed.

### Membrane curvature analysis

Membrane bilayers display elastic characteristics, which resist mechanical bending and stretching. While stretching motions are harder to accomplish, bending motions occur due to thermal fluctuations and/or interactions, e.g., with binding partners. The energetic cost *E* of bending a membrane patch is described by the Helfrich Hamiltonian (Helfrich, 1973; Hu et al., 2012).

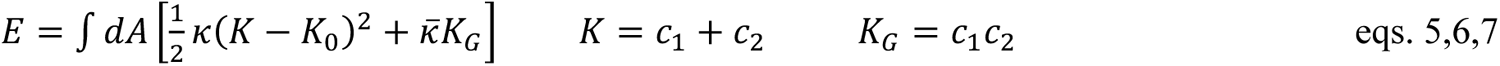

where *c*_1_ and *c*_2_ are the main local curvatures, *K* is the total curvature (*K* / 2 is the mean curvature *H*), *K*_G_ is the Gauss curvature, *κ* is the monolayer bending modulus and *k̅* is the monolayer Gauss curvature modulus. *K*_0_ corresponds to the spontaneous curvature, which is 0 for symmetric bilayers. The integral of the Gauss curvature is 0 for a smooth periodic plane due to symmetry and is a constant for compact surfaces (Hu et al., 2012). Both moduli are quantities difficult to obtain (Nagle, 2013) and to our knowledge, not available for a membrane with a composition DOPE:DOPG 3:1. Here, a value of 12 *kT* was used for *κ*, as determined by X-ray diffraction for DOPE monolayers (Chen and Rand, 1998). It has been shown from simulations that -*k̅* / *k* for DOPE is ∼0.92 (Hu et al., 2012) and we took *k̅* = −*k* for simplicity.

Points representing the surface of each leaflet were obtained using Memsurfer (Bhatia et al., 2019), which triangulates surface points (phosphorous atoms) to generate a surface mesh and uses The Visualization Toolkit (VTK) to obtain their curvatures (Schroeder et al., 2018). Due to fluctuations of lipids on the surface and local high curvatures, the surfaces obtained with default options generated regions with curvature artifacts, particularly in the lower leaflet. To tackle this, the scripts were further modified to add two additional smoothing filters of the retrieved surface points using the VTK python interface: a Laplacian smoothing filter (vtkSmoothPolyDataFilter) and a constrained smoothing filter (vtkConstrainedSmoothingFilter), setting as a constraint distance z-score / 10 of the mean curvature at any given point. While the so generated surfaces are smooth, they include local fluctuations characteristic of Brownian motion, which adds noise when calculating the bending energy (eq. 5).

To measure the curvature caused by the protein and remove noise, the surfaces were fitted to a 2D Gaussian (eq. 8):

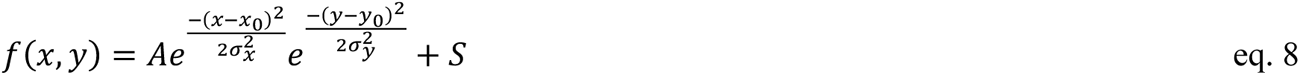

where *x*_0_ and *y*_0_ are the center and *σ*_x_ and *σ*_y_ the variances or spread on the corresponding axis directions, *A* is the amplitude and *S* is a shift term. By expressing *f*(*x*,*y*) implicitly as *F*(*x*,*y*,*z*) = 0, the mean curvature per unit area was calculated (Megrabov, 2014) (eq. 9)

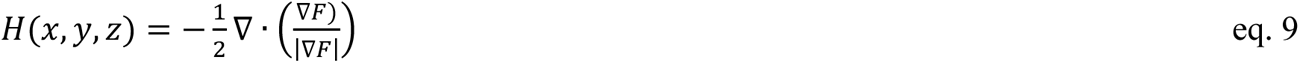

which is the divergence of the normal unit vector to the surface defined by *F*. The Gauss curvature is calculated from the following expression (Trott, 2004)

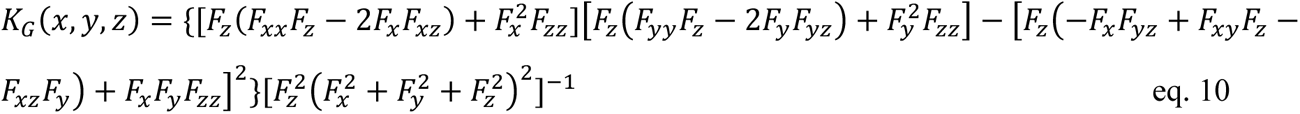

where the subscript denotes a partial derivative. The Gauss curvature term was only calculated for partial surface integrals, as the total for periodic surfaces is 0, as already mentioned (**Fig. 5E**; note the plateau to 0 at higher integration areas).

### Electron cryo-microscopy

PspA grids were prepared by applying 3.5 μL PspA (**Suppl. Table 1**) to glow-discharged (PELCO easiGlow Glow Discharger, Ted Pella Inc.) Quantifoil grids (R1.2/1.3 Cu 200 mesh, Electron Microscopy Sciences). The grids were plunge-frozen in liquid ethane using a ThermoFisher Scientific Vitrobot Mark IV set to 90% humidity at 10 °C (blotting force -5, blotting time 3 to 3.5 s). Movies were recorded in underfocus on a 200 kV Talos Arctica G2 (ThermoFisher Scientific) electron microscope equipped with a Bioquantum K3 (Gatan) detector operated by EPU (ThermoFisher Scientific).

Tilt series for Cryo-ET were recorded on a 300 kV Titan Krios G4 (ThermoFisher Scientific) electron microscope equipped with a Biocontinuum K3 (Gatan) detector operated by Tomo (ThermoFisher Scientific) using a dose-symmetric scheme at -60° to 60° with 3° steps. Tilt images were acquired at a magnification of 64 kx (pixel size 1.362 Å) with a nominal underfocus of 4.0 µm. The total dose for each tilt series was 131 e^−^/Å^2^. A total of 20 tilt series were collected.

### Cryo-electron tomography image processing

Tilt image frames were gain-corrected, dose-weighted, and aligned using WARP (Tegunov and Cramer, 2019). The resulting tilt series were aligned, 8x binned, and reconstructed by the weighted back projection method using AreTomo (Zheng et al., 2022). The tomograms were segmented in Dragonfly (Object Research Systems) by progressively training a U-Net with an increasing number of manually segmented tomogram frames (5-15). Then the trained U-Net was used to predict those features of the tomogram. For visualization, the segmentation was cleaned up by removing isolated voxels of each label group (islands of <150 unconnected voxels). The resulting segmentation was imported to ChimeraX (Pettersen et al., 2021) for 3D rendering.

### Image processing and helical reconstruction

Movie frames were gain corrected, dose weighted, and aligned using cryoSPARC Live (Punjani et al., 2017). Initial 2D classes were produced using the auto picker implemented in cryoSPARC Live. The following image processing steps were performed using cryoSPARC. The best-looking classes were used as templates for the filament trace. The resulting filament segments were extracted with 600 px box size (Fourier cropped to 200 px) and subjected to multiple rounds of 2D classification. The remaining segments were reextracted with a box size of 400 px (Fourier cropped to 200 px) and subjected to an additional round of 2D classification. The resulting 2D class averages were used to determine filament diameters and initial symmetry guesses in PyHI (Zhang, 2022). Symmetry guesses were validated by initial helical refinement in cryoSPARC and selection of the helical symmetry parameters yielding reconstructions with typical PspA features and the best resolution. Then all segments were classified by heterogeneous refinement and subsequent 3D classifications using the initial helical reconstructions as templates. The resulting class distribution gave the PspA rod diameter distribution shown in **Figure 3**. The resulting helical reconstructions were subjected to multiple rounds of helical refinement including the symmetry search option. For the final polishing, the segments were reextracted at 400 px without Fourier cropping. Bad segments were discarded by heterogeneous refinement. Higher-order aberrations were corrected by global and local CTF refinement followed by a final helical refinement step. The local resolution distribution and local filtering was performed using cryoSPARC (**Suppl. Fig. 1A-C**). The resolution of the final reconstructions was determined by Fourier shell correlation (auto-masked, FSC=0.143) (**Suppl. Fig. 1D**).

### Cryo-EM map interpretation and model building

The 3D reconstructions were B-factor sharpened in Phenix (*phenix.auto-sharpen*) (Terwilliger et al., 2018). The handedness of the final map was determined by rigid-body fitting the PspA reference structure aa 22-217 (PDB:7ABK) (Junglas et al., 2021) into the final maps using ChimeraX (Goddard et al., 2018; Pettersen et al., 2021) and flipped accordingly. 7ABK was flexibly MDFF fitted to the 3D reconstructions using ISOLDE (Croll, 2018). Some of the structures in the presence of EPL showed additional density at the tip of α1 which we interpreted as the additional N-terminal residues. Therefore, helix α0 (aa 1-22) was built from scratch, joined with α1 and flexibly MDFF fitted to the 3D reconstructions with helix restraints from aa 2-9 and aa 11-21 using ISOLDE (Croll, 2018). Then the respective helical symmetry was applied to all models to create assemblies of 60 monomers each. The assembly models were subjected to auto-refinement with *phenix.real_space_refine* (Afonine et al., 2018b) (with NCS constraints and NCS refinement). After auto-refinement, the new models were used for local model-based map sharpening with LocSCALE (Jakobi et al., 2017) to produce the final maps. The auto-refined models were checked/adjusted manually in Coot (Emsley et al., 2010) and ISOLDE (Croll, 2018) before a final cycle of auto-refinement with *phenix.real_space_refine* (Afonine et al., 2018b) (with NCS constraints and NCS refinement). After the final inspection, the model was validated in *phenix.validation_cryoem (Afonine et al., 2018a)/Molprobity (Williams et al., 2018).* Image processing, helical reconstruction, and model building were completed using SBGrid-supported applications (Morin et al., 2013). In this manner, a total of 15 cryo-EM maps were determined in 2 samples (see **Tables 1 and2**).

**Table 1.**
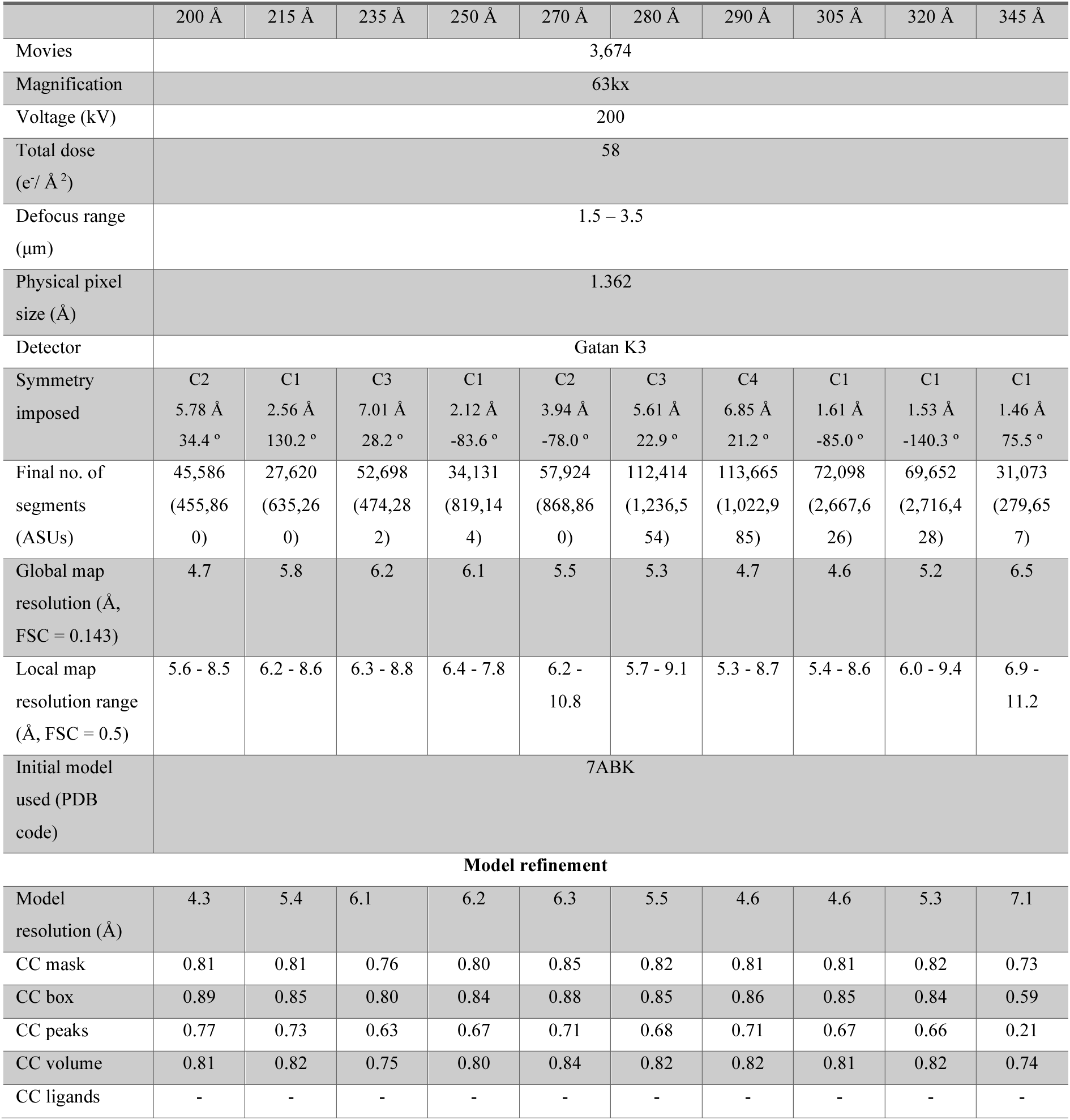
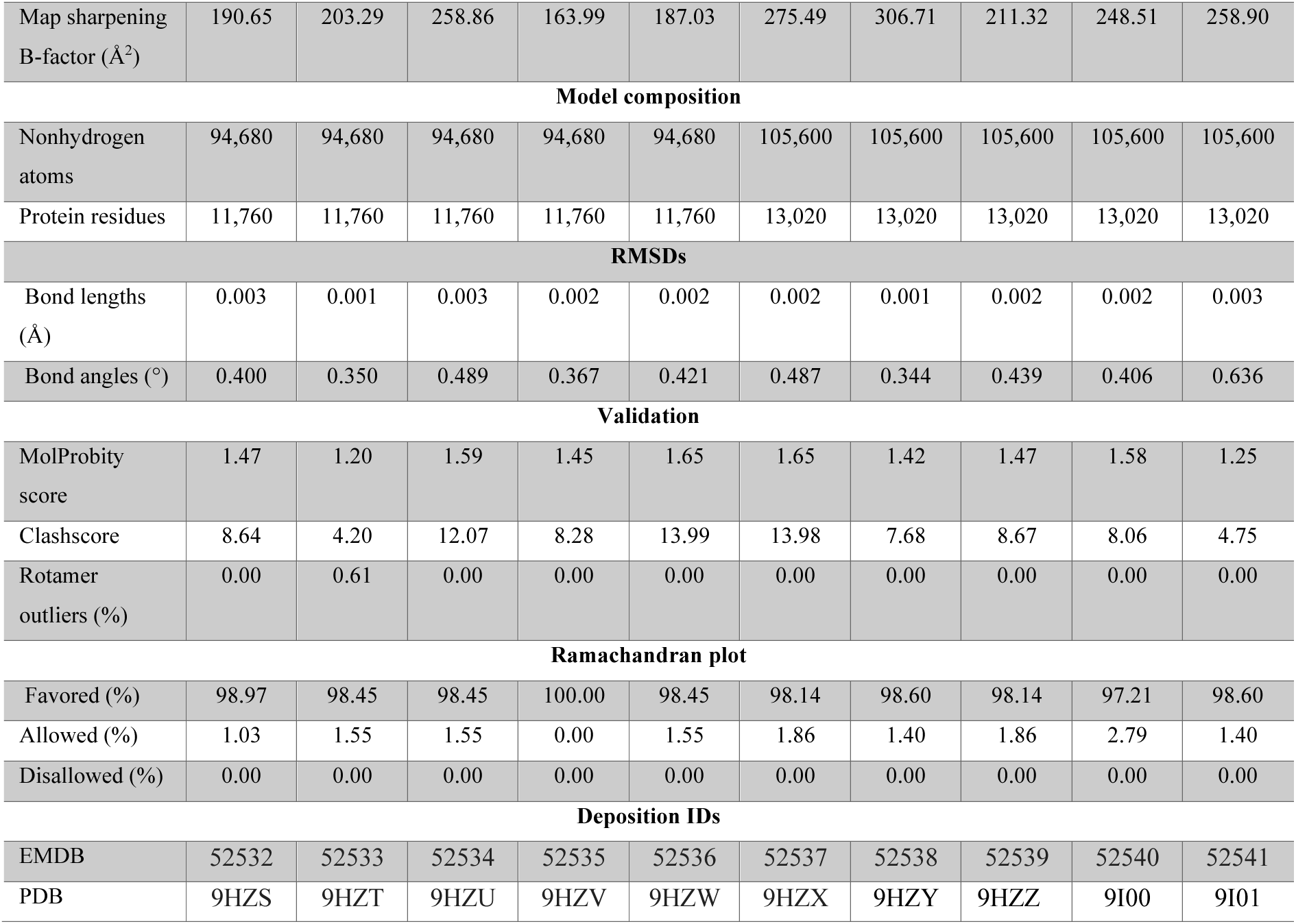
Data collection, image processing, and model refinement PspA EPL.

**Table 2.** Data collection, image processing, and model refinement PspA α1-5 EPL.

| | 235 $\text{\AA}$ | 250 $\text{\AA}$ | 270 $\text{\AA}$<br>C1 | 270 $\text{\AA}$<br>C2 | 280 $\text{\AA}$ | 290 $\text{\AA}$ |
| --- | --- | --- | --- | --- | --- | --- |
| Movies | 3,149 |  |  |  |  |  |
| Magnification | 49kx |  |  |  |  |  |
| Voltage (kV) | 200 |  |  |  |  |  |
| Total dose<br>($e^-/\text{\AA}^2$) | 44 | | | | | |
| Defocus range<br>($\mu\text{m}$) | 1.0 – 3.5 | | | | | |
| Physical pixel<br>size ($\text{\AA}$) | 0.868 | | | | | |
| Detector | Gatan K3 |  |  |  |  |  |
| Symmetry<br>imposed | C4<br>9.03 $\text{\AA}$<br>27.6 $^\circ$ | C1<br>2.10 $\text{\AA}$<br>-83.6 $^\circ$ | C1<br>1.92 $\text{\AA}$<br>76.7 $^\circ$ | C2<br>3.93 $\text{\AA}$<br>-78.0 $^\circ$ | C5<br>9.03 $\text{\AA}$<br>21.9 $^\circ$ | C4<br>6.85 $\text{\AA}$<br>21.1 $^\circ$ |
| Final no. of<br>segments<br>(ASUs) | 121,749<br>(852,243) | 1,405,435<br>(19,676,090) | 629,133<br>(10,066,128) | 568,968<br>(4,551,744) | 636,672<br>(1,910,016) | 166,757<br>(1,500,813) |
| Global map<br>resolution ($\text{\AA}$,<br>FSC = 0.143) | 4.2 | 3.8 | 3.9 | 3.9 | 3.9 | 5.4 |
| Local map resolution range (Å, FSC = 0.5) | 4.4 - 5.1 | 3.5 - 4.8 | 3.9 - 5.1 | 3.9 - 5.2 | 4.0 - 5.2 | 5.5 - 6.4 |
| Initial model used (PDB code) | 7ABK |  |  |  |  |  |
| Model refinement |  |  |  |  |  |  |
| Model resolution (Å) | 4.1 | 3.7 | 3.8 | 3.9 | 3.8 | 5.0 |
| CC mask | 0.81 | 0.77 | 0.74 | 0.76 | 0.79 | 0.75 |
| CC box | 0.88 | 0.84 | 0.82 | 0.83 | 0.86 | 0.83 |
| CC peaks | 0.77 | 0.70 | 0.670 | 0.72 | 0.74 | 0.66 |
| CC volume | 0.81 | 0.76 | 0.74 | 0.76 | 0.79 | 0.76 |
| Map sharpening B-factor (Å²) | 147.92 | 29.07 | 97.75 | 115.88 | 128.22 | 240.89 |
| Model composition |  |  |  |  |  |  |
| Nonhydrogen atoms | 93,600 | 93,600 | 93,600 | 93,600 | 93,600 | 93,600 |
| Protein residues | 11,640 | 11,640 | 11,640 | 11,640 | 11,640 | 11,640 |
| RMSDs |  |  |  |  |  |  |
| Bond lengths (Å) | 0.002 | 0.002 | 0.003 | 0.004 | 0.002 | 0.003 |
| Bond angles (°) | 0.374 | 0.480 | 0.643 | 0.881 | 0.436 | 0.477 |
| Validation |  |  |  |  |  |  |
| MolProbity score | 1.44 | 1.54 | 1.53 | 1.36 | 1.40 | 1.53 |
| Clashscore | 8.03 | 10.42 | 10.21 | 6.50 | 6.97 | 10.11 |
| Rotamer outliers (%) | 0.00 | 0.31 | 0.00 | 0.75 | 0.62 | 0.62 |
| Ramachandran plot |  |  |  |  |  |  |
| Favored (%) | 98.96 | 98.30 | 98.44 | 100.00 | 97.92 | 98.44 |
| Allowed (%) | 1.04 | 1.70 | 1.56 | 0.00 | 2.08 | 1.56 |
| Disallowed (%) | 0.00 | 0.00 | 0.00 | 0.00 | 0.00 | 0.00 |
| Deposition IDs |  |  |  |  |  |  |
| EMDB | 52526 | 52527 | 52528 | 52529 | 52530 | 52531 |
| PDB | 9HZM | 9HZN | 9HZO | 9HZP | 9HZQ | 9HZR |

Diameters of the final PspA rod reconstructions were measured using ImageJ (Rueden et al., 2017): First radial intensity profiles of each reconstruction were created (**Suppl. Fig. 1A**). Then the radius at an intensity cutoff of 0.3 was read. The radius readout from reconstructions from the same helical symmetry class was averaged, converted to diameters and rounded to 5 Å increments. Outer and inner leaflet radii of engulfed membrane tubes were determined from the same radial intensity profiles at the peak maxima from each bilayer leaflet.

### Analysis and visualization tools

Metrics obtained from molecular simulations and trajectories were handled with CPPTRAJ (Roe and Cheatham, 2013). Visualization of structures was done using Open-Source PyMOL version 2.4 (DeLano, 2002), ChimeraX 1.6.1 (Pettersen et al., 2021) and VMD 1.9.4 VM (Humphrey et al., 1996). The fitting process and all calculations were done using Python 3 and NumPy (Harris et al., 2020), SciPy (Virtanen et al., 2020), and SymPy (Meurer et al., 2017). Data and statistical analysis were performed using OriginPro 2021b (OriginLab Corp., Northampton, USA). Detailed descriptions of quantifications and statistical analyses (exact values of *n*, dispersion and precision measures used, and statistical tests used) can be found in the respective Figures, Figure legends, and Methods section.

## Author contributions

E.H., B.J., S.S.V., D.S., H.G., and C.S. designed the research. E.H. and I.R. cloned, expressed, and purified the proteins. E.H. and B.J. prepared cryo-EM samples. E.H. and B.J. operated the electron microscopes. E.H., B.J., and C.S. determined the cryo-EM structures. E.H. and B.J. built the refined atomic models. M.K. performed the membrane binding assays. M.K., N.H., and D.S. analyzed the membrane binding data and wrote the respective parts. S.S.V. performed MD simulations. S.S.V. and H.G. analyzed MD simulations and wrote the respective parts. E.H., B.J., S.S.V, D.S., H.G., and C.S. wrote the manuscript with input from all authors.

## Declaration of interests

The authors declare no competing interests.

## Materials availability

All unique and stable reagents generated in this study are available from the Lead Contact with a completed Material Transfer Agreement.

## Data and code availability

The EMDB accession numbers for cryo-EM maps and PspA models are EMD IDs: 52526, 52527, 52528, 52529, 52530, 52531, 52532, 52533, 52534, 52535, 52536, 52537, 52538, 52539, 52540, and 52541 and PDB-IDs: 9HZM, 9HZN, 9HZO, 9HZP, 9HZQ, 9HZR, 9HZS, 9HZT, 9HZU, 9HZV, 9HZW, 9HZX, 9HZY, 9HZZ, 9I00, and 9I01.

## Acknowledgments

This study was funded by the Deutsche Forschungsgemeinschaft (DFG, German Research Foundation, SA 1882/6-1, SCHN 690/16-1, SFB1551 (Project number 464588647), SFB1552 (Project number 465145163) and SFB1208 (Project number 267205415)). The authors gratefully acknowledge the electron microscopy access time and computing time granted by the biological EM facility of the Ernst-Ruska Centre at Forschungszentrum Jülich. In this regard, we thank Thomas Heidler and Pia Sundermeyer for maintaining the electron microscopes and Daniel Mann for maintaining the processing computers. The authors gratefully acknowledge the computing time granted through Jülich Aachen Research Alliance (JARA) on the supercomputer JURECA at Forschungszentrum Jülich (Krause, 2019), the computational support and infrastructure provided by the “Zentrum für Informations-und Medientechnologie” (ZIM) at the Heinrich Heine University Düsseldorf, and the computing time provided by the John von Neumann Institute for Computing (NIC) on the supercomputer JUWELS at Jülich Supercomputing Centre (JSC) (user ID: VSK33).

## Supplementary Information

**Suppl. Table 1.**
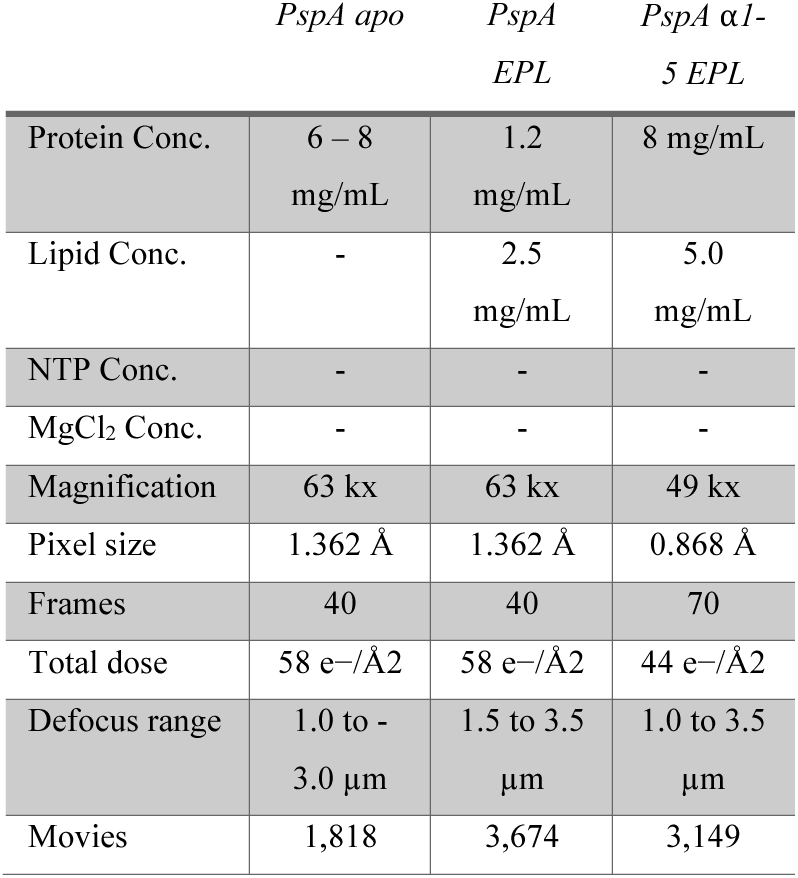
Sample Details.

**Suppl. Table 2.** Bessel order overview of SynPspA + EPL.

|  | 200 Å | 215 Å | 235 Å | 250 Å | 270 Å | 280 Å | 290 Å | 305 Å | 320 Å | 345 Å |
| --- | --- | --- | --- | --- | --- | --- | --- | --- | --- | --- |
| **Bessel order 1.** | 10 | 11 | 12 | 13 | 14 | 15 | 16 | 17 | 18 | 19 |
| Layer Line<br>position (1/ Å) | 121.1 | 92.3 | 118.4 | 109.0 | 121.1 | 115.9 | 115.9 | 113.5 | 113.5 | 113.5 |
| **Bessel order 2.** | 2 | 3 | 3 | 4 | 4 | 3 | 4 | 4 | 5 | 5 |
| Layer Line<br>position (1/ Å) | 30.3 | 30.8 | 29.6 | 30.3 | 30.3 | 29.0 | 29.0 | 28.4 | 28.4 | 28.4 |

**Suppl. Figure 1:**
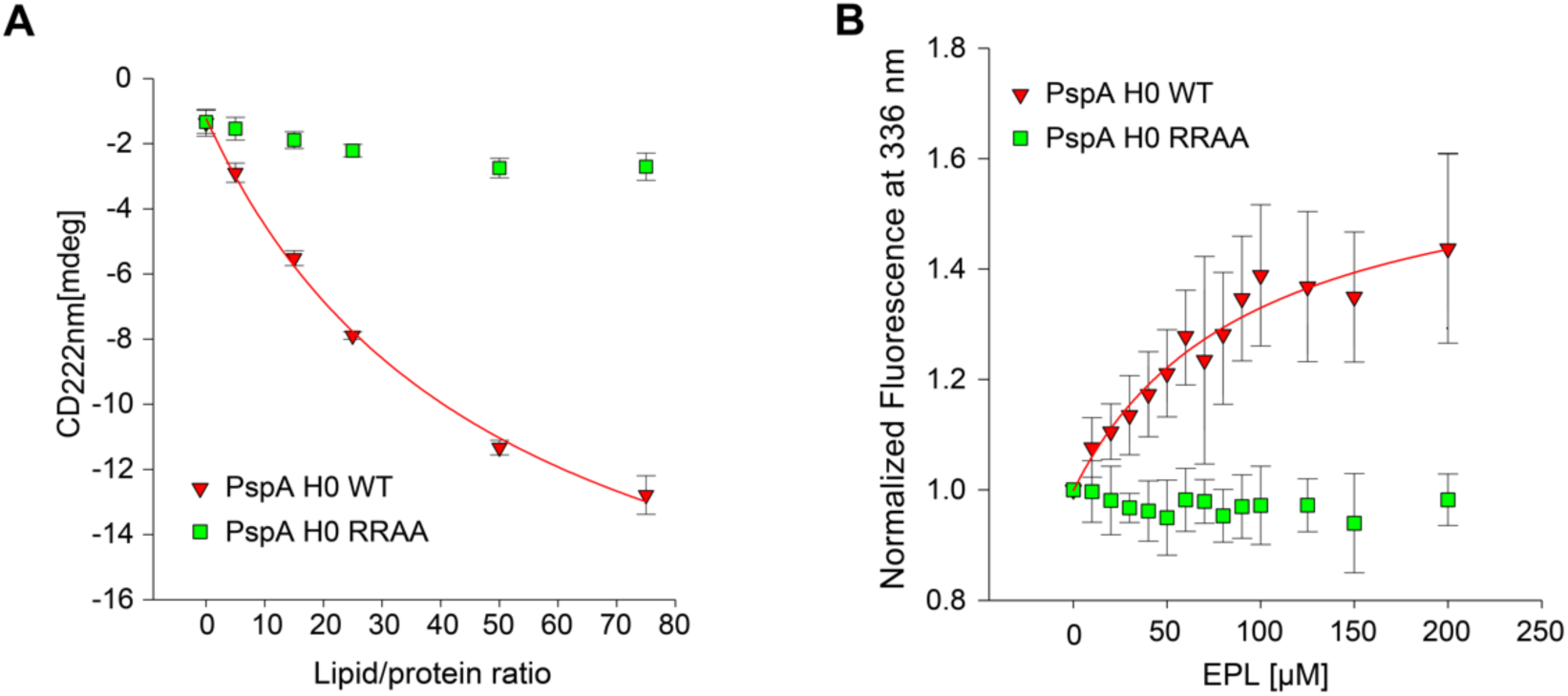
Interaction of PspA’s isolated helix α0 with membranes. **A:** Changes in ellipticity at 222 nm based on the circular dichroism (CD) of 34 µM helix α0 peptides at increasing EPL/peptide ratios spectra shown in Figure 1B, green: PspA wild-type (WT), red: PspA α0 RRAA mutant) indicate an EPL-induced increase in α-helix content in the WT peptide. The binding data obtained for the WT peptide by plotting the ellipticity at 222 nm versus the EPL/peptide ratio were analyzed based on a hyperbolic function yielding an apparent partition coefficient of 49.8 ± 8.2. **B:** Changes in the Trp-fluorescence of 2 µM helix α0 in the presence of EPL at increasing EPL/peptide ratios reveal a fluorescence emission maximum change for the WT peptide but no shift for the PspA α0 RRAA (Fig. 1). The binding data obtained by plotting the fluorescence intensities at 336 nm versus EPL/peptide ratio were analyzed based on a hyperbolic function, yielding an apparent partition coefficient of 47.0 ± 15.4. Thus, a lipid/peptide ratio similar to the value determined by CD spectroscopy is required to bind 50 % of the peptide. Based on these results, we estimated the free energy of binding of the WT peptide from the CD spectroscopy and Trp-based fluorescence analyses to be between Δ*G*^0^ = -3.9 and -3.7 kcal mol^-1^, and -5.7 and -5.3 kcal mol^-1^, respectively.

**Suppl. Figure 2:**
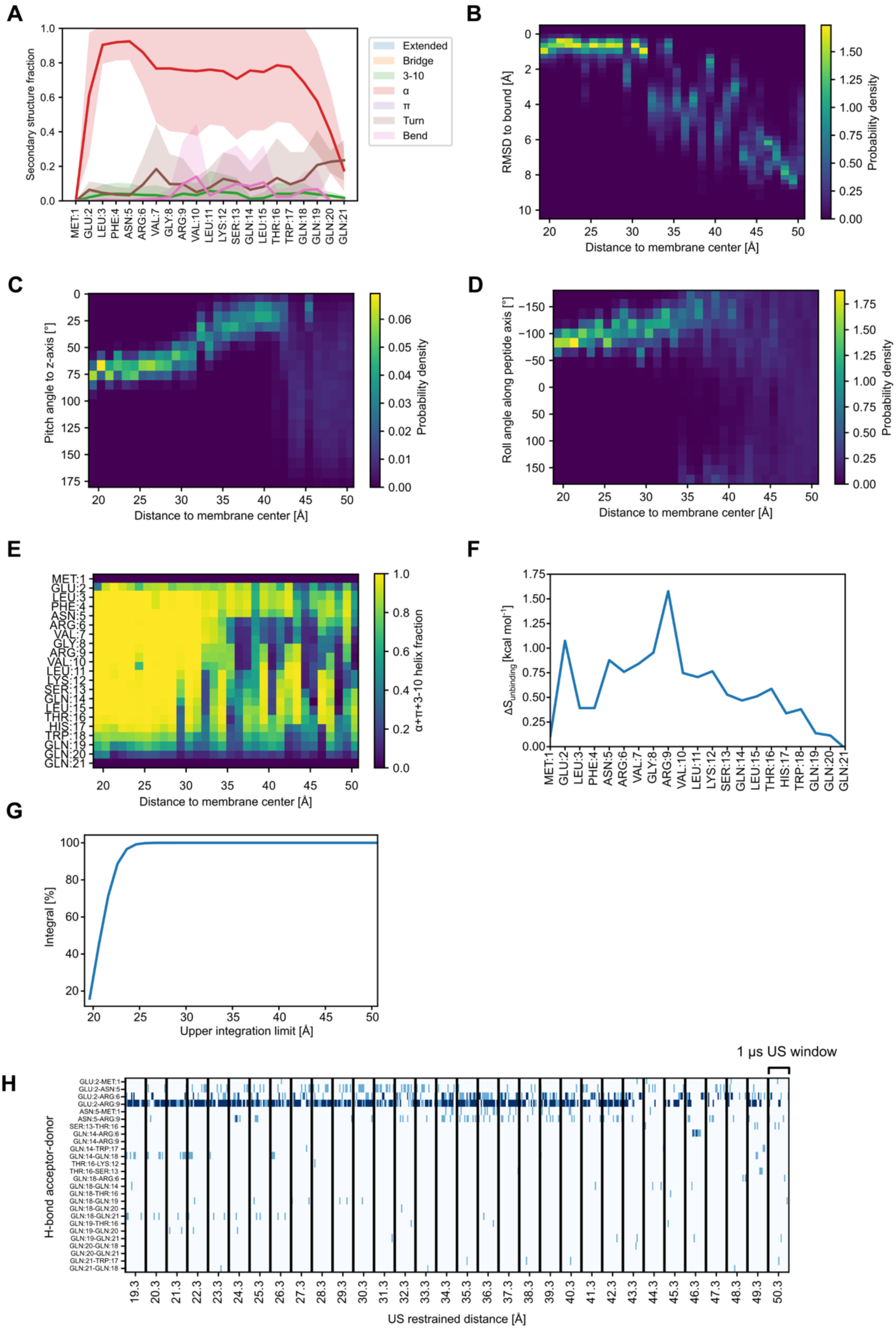
Helix α0 metrics in unbiased molecular dynamics (MD) and umbrella sampling (US) simulations. **A**: Residue-wise mean secondary structure content over 12 replicas of unbiased MD simulations (as calculated by cpptraj) with the shaded area showing the standard deviation. On average, α0 forms an α-helix upon binding to the membrane surface. **B-E**: Analysis of peptide orientation and structural changes in the 32 umbrella sampling windows, with the reaction coordinate representing the distance to the membrane center along the z-axis (membrane normal). Along the reaction coordinate, the global degrees of freedom of helix α0 are restricted, as indicated by (**B**) the RMSD of the sampled structures, (**C**) the Euler angle describing the pitch with respect to the membrane normal, and (**D**) the roll angle describing how much the peptide can roll around the peptide axis. **E**: As the peptide unbinds, the peptide transitions from a fully helical structure at the membrane surface to a partially unstructured conformation, with partial helical sections towards the N and C terminal ends. **F**: PDB2ENTROPY allows one to estimate conformational entropy changes between structure ensembles by performing a *k*-nearest neighbors analysis of the torsions of the peptide. **G**: The integration of the PMF (integral term in eq. 4) yields an almost constant value for an upper integration limit beyond 25 Å. **H**: Hydrogen bonds between side chains with a cumulative prevalence of at least 100 ns (10% of a single window). Light blue depicts a single H-bond, while dark blue shows that two H-bonds are involved in the interaction.

**Suppl. Figure 3:**
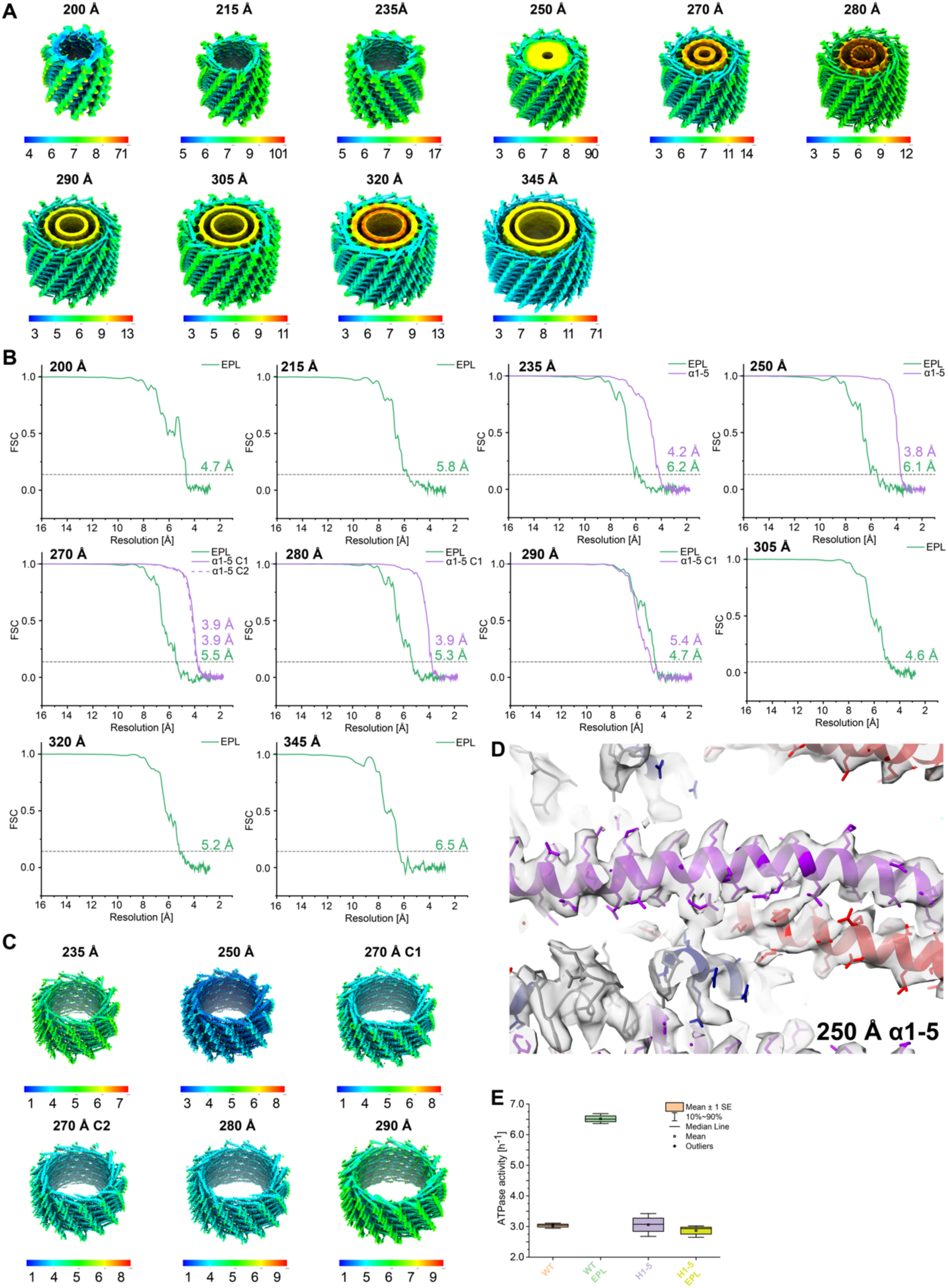
Local Resolution maps and FSC curves of cryo-EM PspA structures. **A**: Local resolution maps of the helical PspA assemblies of the PspA + EPL sample with resolution displayed on the isosurface (scale indicated below). **B**: FSC curves of all PspA diameters of the data sets with PspA+EPL (green) and the data set with PspA *α*1-5 + EPL (violet) including the 0.143 resolution dashed threshold line. **C:** Local resolution maps of the helical PspA *α*1-5 + EPL assemblies with resolution displayed on isosurface (scale indicated below). **D:** 3.8 Å resolution density of 250 Å wide PspA *α*1-5 + EPL rod assembly with the refined atomic model of the monomer in ribbon representation (*α*1 red, *α*2+3 violet, *α*4 blue, *α*5 cyan). **E**: ATPase activity of the wild-type protein and the *α*1-5 mutant in the absence and presence of EPL; wild-type (WT) (orange): ATPase activity without EPL; WT EPL (green): ATPase activity in the presence of EPL; *α*1-5 variant (purple): ATPase activity of *α*1-5 variant in the absence of EPL; *α*1-5 EPL (yellow): ATPase activity of *α*1-5 variant in the absence of EPL; n=3.

**Suppl. Figure 4:**
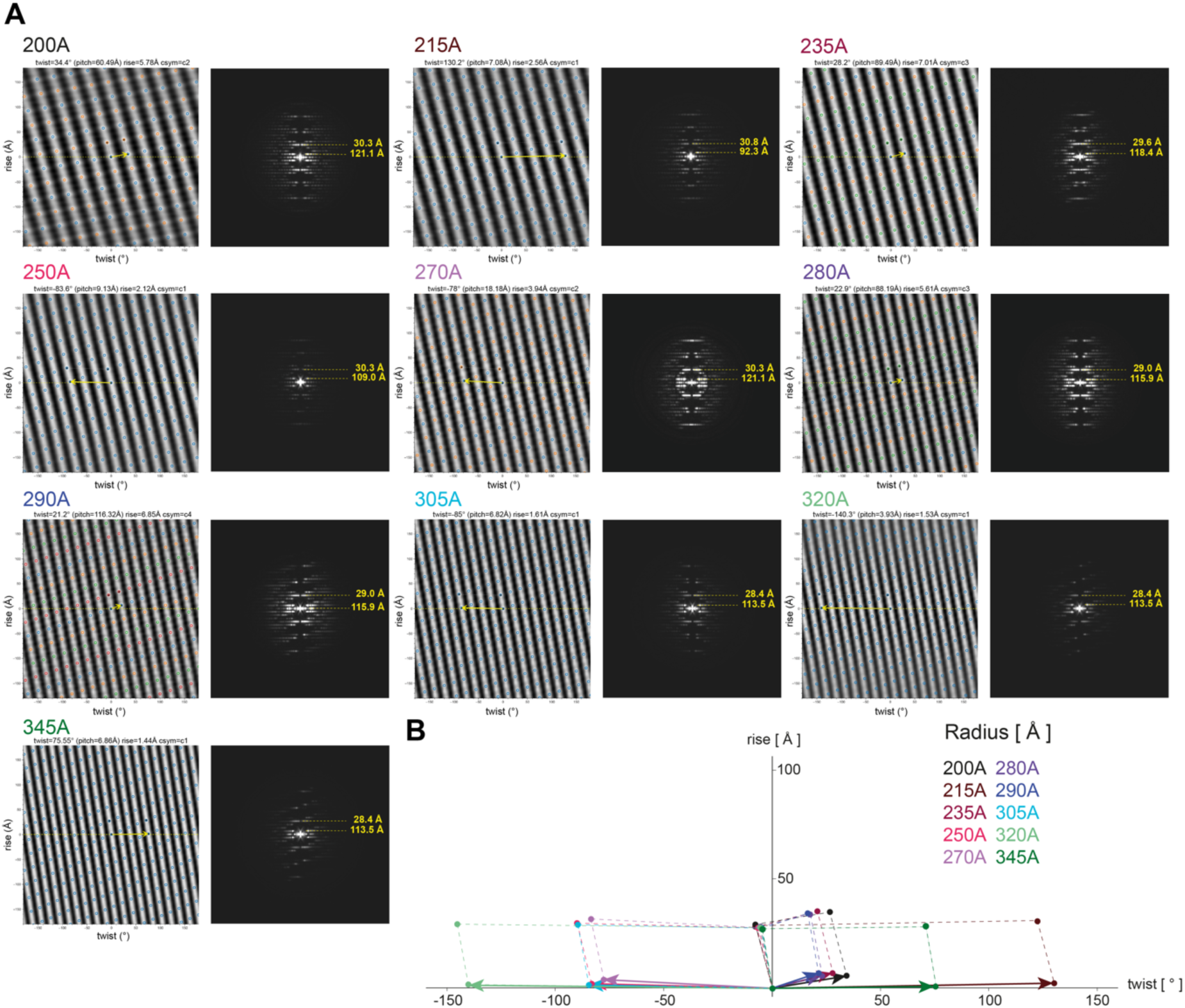
Helical lattice and Power spectra of PspA + EPL structures. **A**: Helix lattice plot (left) generated with HI3D (Sun et al., 2022), including unit cell of symmetry vectors and power spectra (right) generated using PyHI (Zhang, 2022) together with used layer line position (yellow dashed lines) for PspA + EPL structure determination. **B**: Superimposed unit cells of the symmetry vectors of the different PspA diameters.

**Suppl. Figure 5:**
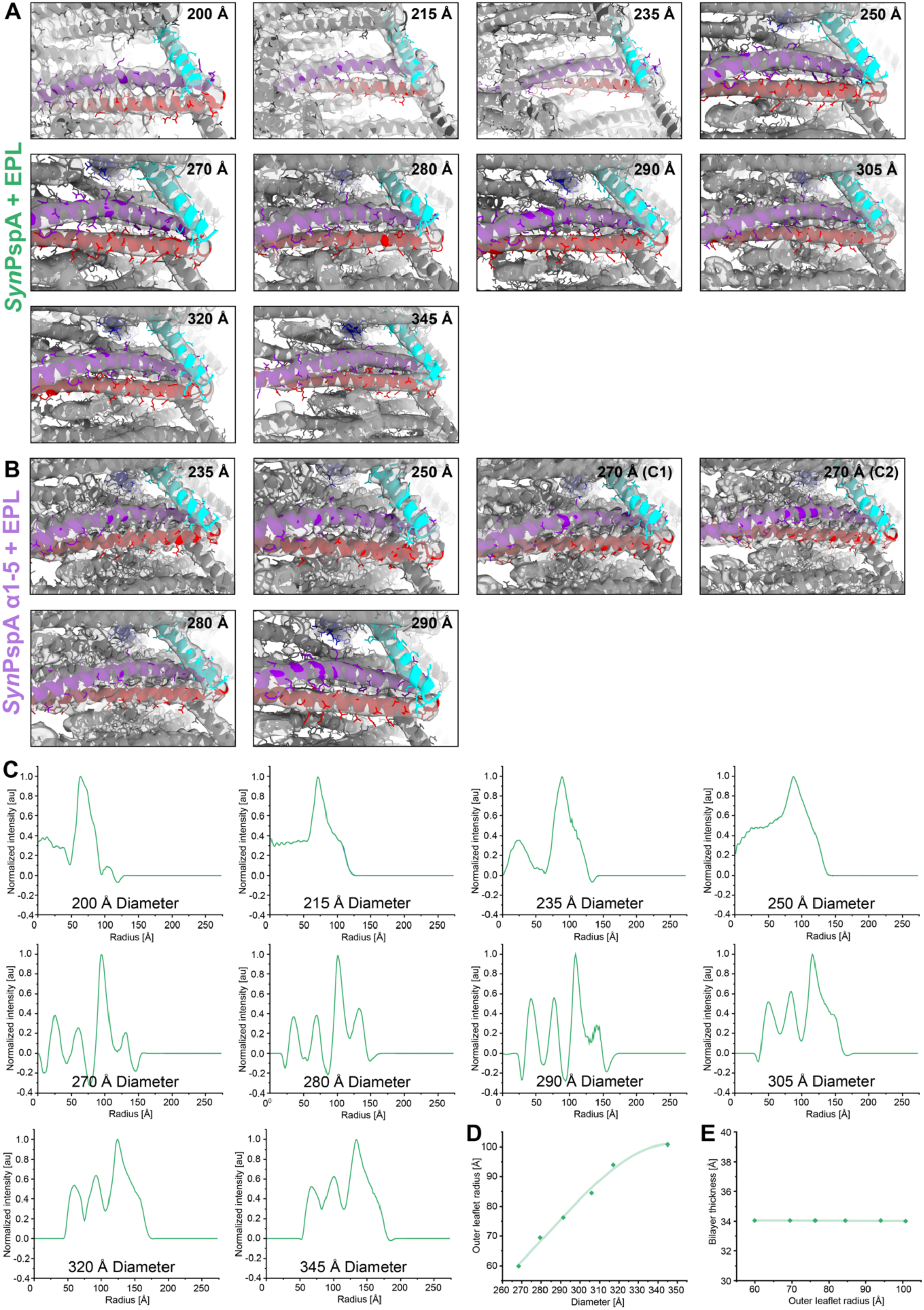
PspA rod density, radial density profiles, and mass per length. **A:** Density of PspA+EPL assemblies of different diameter rods with the atomic model of the monomer in ribbon representation (*α*1 red, *α*2+3 violet, *α*4 blue, *α*5 cyan). **B**: Density of PspA *α*1-5 +EPL assemblies of different diameter rods with the atomic model of the monomer in ribbon representation (*α*1 red, *α*2+3 violet, *α*4 blue, *α*5 cyan) **C**: Radial density profiles of PspA+EPL rods with diameters ranging from 200 - 345 Å showing typical peaks for a lipid bilayer and the protein density. **D:** Scatter plot of the outer leaflet membrane tube radius from PspA+EPL rods, measured as the max/min from the density profile against the respective bilayer thickness. **E:** Scatter plot of PspA+EPL rods diameters containing significant tubular membrane density against the respective outer leaflet radius of the membrane tube.

**Suppl. Figure 6:**
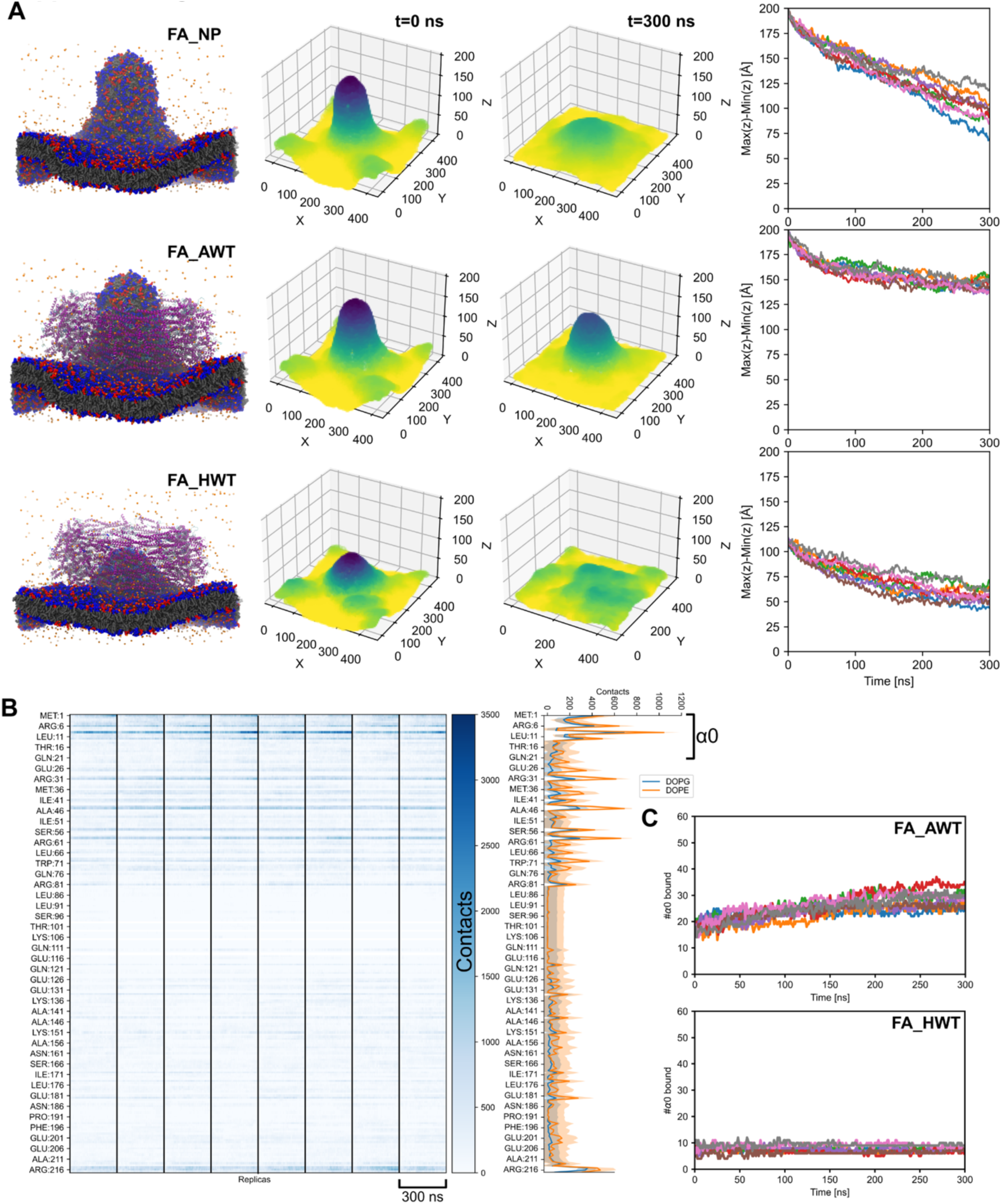
The 290 Å PspA bilayer complex modeled in the full-atom representation. **A**: Left: Depiction of full-atom systems in the absence (FA_NP, top) and the presence of PspA, with the curved membrane spanning fully (FA_AWT, middle) and halfway through (FA_HWT, bottom). Center: Surface depictions of the initial (*t* = 0 ns) and final (*t* = 300 ns) structures of the upper leaflet of the membrane. Right: The distance between the highest and lowest point of the membrane along the membrane normal as a proxy of the change in membrane curvature. **B**: Summed per-residue number of contacts over the 60 simulated PspA chains in the 290 Å complex with the lipid headgroups on the membrane surface, over eight replicas. To the right, the mean number of contacts with DOPG and DOPE lipids along the trajectory is given, with the standard deviation as the shaded area. **C**: Total number of helices α0 interacting with the membrane during the simulations for FA_AWT (top) and FA_HWT (bottom).

**Suppl. Figure 7:**
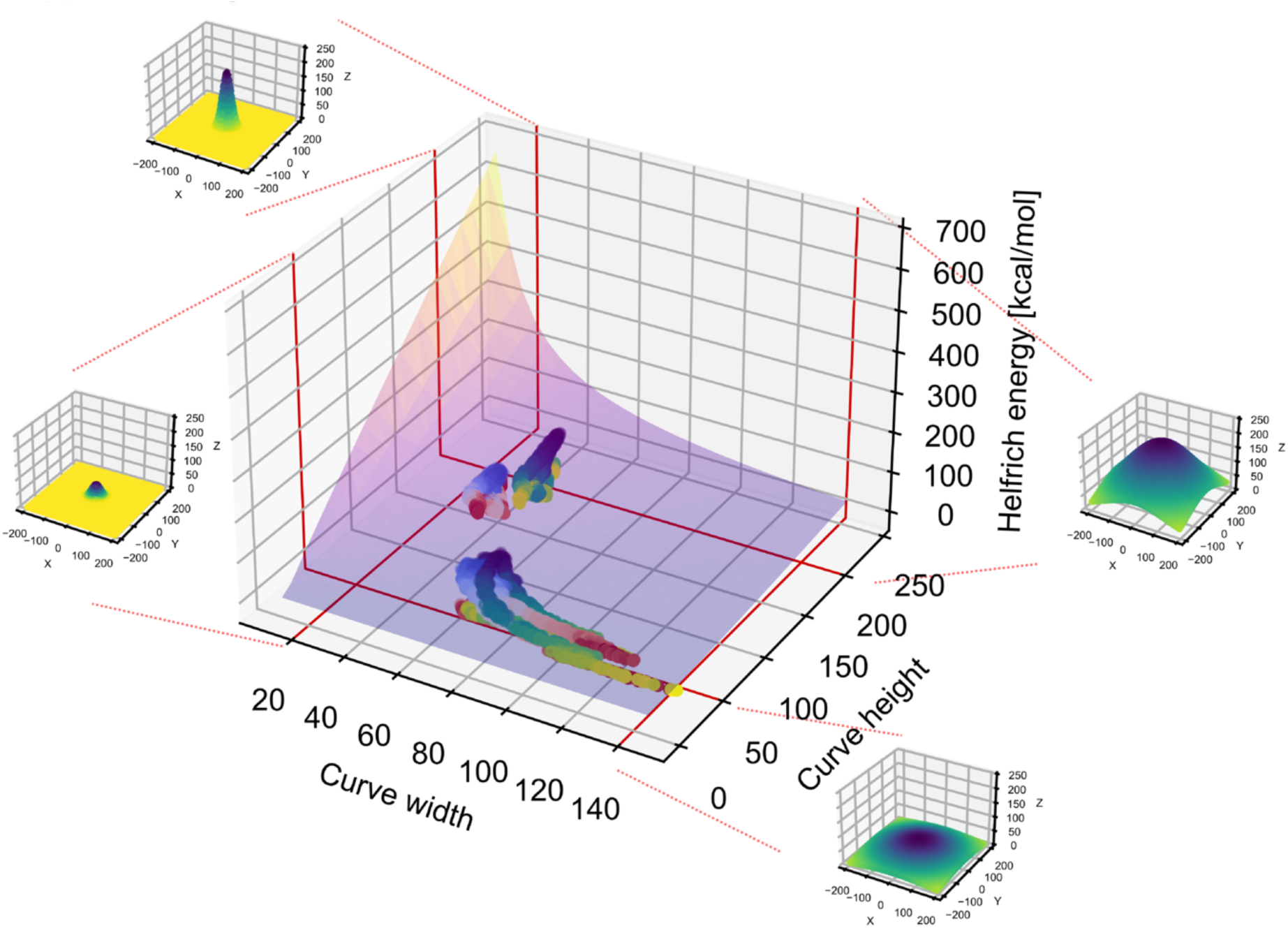
Helfrich bending energy for 2D Gaussians of different amplitudes and standard deviations. The depicted graph surface is the same as that shown in **Fig. 5D**, here given as a 3D surface rather than a contour plot. Additional Gaussian surfaces for low/high width and height are included as visual aids. Helfrich energies of snapshots of full-atom all-way-through (FA_AWT) and full-atom half-way-through (FA_HWT) trajectories are projected onto the energy surface (blue to yellow, for start to finish upper leaflet, and light blue to red, for start to finish lower leaflet). For further details, refer to the caption of **Fig. 5D**

**Suppl. Figure 8:**
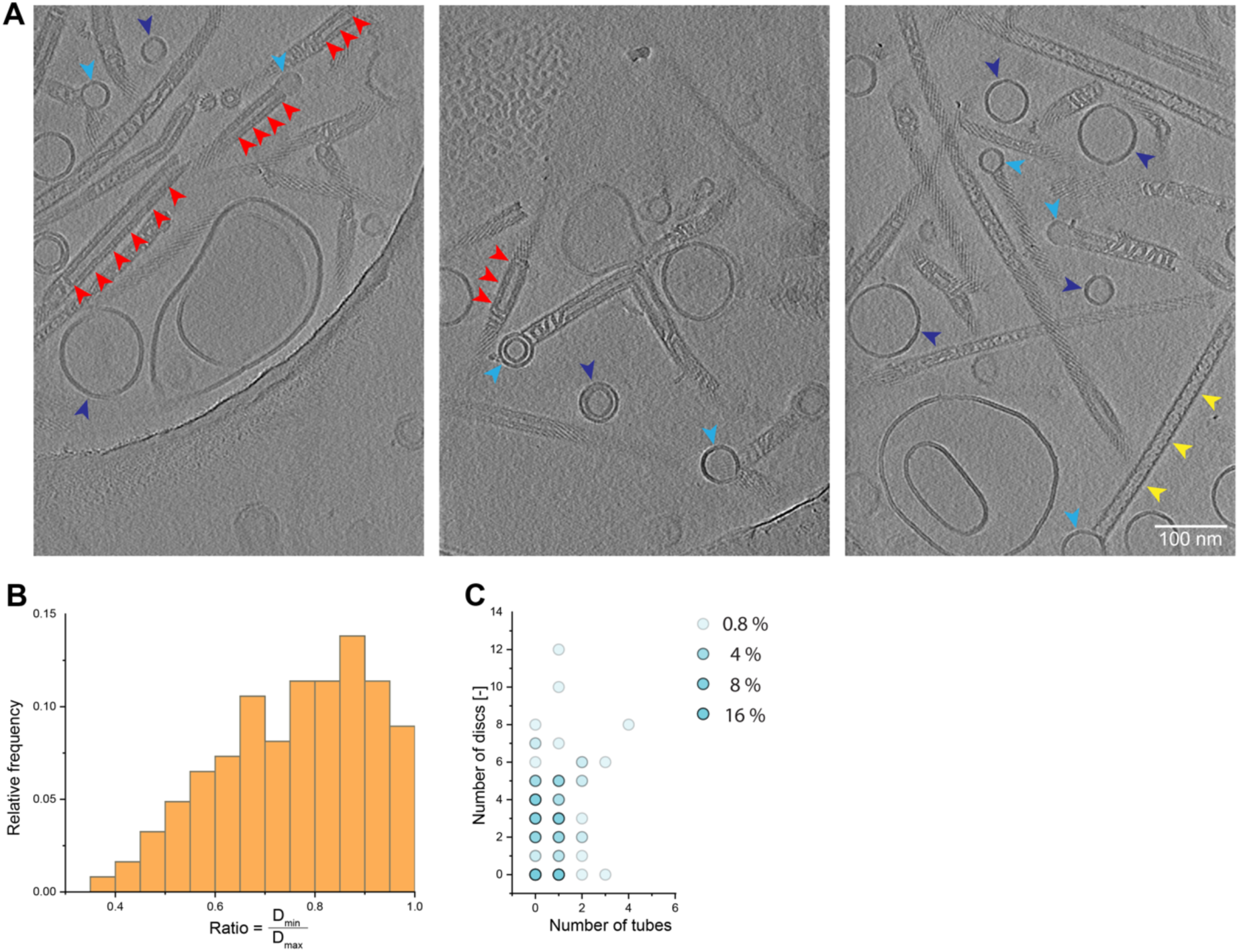
Tomograms of PspA with EPL membranes. **A**: Tomogram images of PspA with EPL membranes. Red arrows: completely tubulated EPL vesicles within the PspA rod structures, dark blue arrows: stand-alone vesicles, light blue arrows: rod-attached vesicles, red arrows: rod-internalized vesicles, yellow arrows: PspA rod without EPL membrane in the rod lumen. **B**: Histogram of the diameter variation per rod, plotted as the ratio of the minimum and maximum diameter within a rod (n = 125). **C**: Dependency distribution of the membrane structures within the rods shown as a scatter plot (n = 125).

## Notes

### Competing Interest Statement

The authors have declared no competing interest.

### Summary of Updates

Title had a typo included. Updated Title: "The bacterial ESCRT-III PspA rods thin lipid tubules and increase membrane curvature through helix α0 interactions"

## References

Afonine, P.V., Klaholz, B.P., Moriarty, N.W., Poon, B.K., Sobolev, O.V., Terwilliger, T.C., Adams, P.D., Urzhumtsev, A., 2018a. New tools for the analysis and validation of cryo-EM maps and atomic models. Acta Cryst D 74, 814–840. 10.1107/S2059798318009324

Afonine, P.V., Poon, B.K., Read, R.J., Sobolev, O.V., Terwilliger, T.C., Urzhumtsev, A., Adams, P.D., 2018b. Real-space refinement in PHENIX for cryo-EM and crystallography. Acta Cryst D 74, 531–544. 10.1107/S2059798318006551

Ahmad, S., Strunk, C.H., Schott-Verdugo, S.N., Jaeger, K.-E., Kovacic, F., Gohlke, H., 2021. Substrate Access Mechanism in a Novel Membrane-Bound Phospholipase A of *Pseudomonas aeruginosa* Concordant with Specificity and Regioselectivity. J. Chem. Inf. Model. 61, 5626– 5643. 10.1021/acs.jcim.1c00973

Azad, K., Guilligay, D., Boscheron, C., Maity, S., De Franceschi, N., Sulbaran, G., Effantin, G., Wang, H., Kleman, J.-P., Bassereau, P., Schoehn, G., Roos, W.H., Desfosses, A., Weissenhorn, W., 2023. Structural basis of CHMP2A–CHMP3 ESCRT-III polymer assembly and membrane cleavage. Nat Struct Mol Biol 30, 81–90. 10.1038/s41594-022-00867-8

Barrera, E.E., Machado, M.R., Pantano, S., 2019. Fat SIRAH: Coarse-Grained Phospholipids To Explore Membrane–Protein Dynamics. J. Chem. Theory Comput. 15, 5674–5688. 10.1021/acs.jctc.9b00435

Ben-Shalom, I.Y., Pfeiffer-Marek, S., Baringhaus, K.-H., Gohlke, H., 2017. Efficient Approximation of Ligand Rotational and Translational Entropy Changes upon Binding for Use in MM-PBSA Calculations. J. Chem. Inf. Model. 57, 170–189. 10.1021/acs.jcim.6b00373

Ben-Shaul, A., Ben-Tal, N., Honig, B., 1996. Statistical thermodynamic analysis of peptide and protein insertion into lipid membranes. Biophysical Journal 71, 130–137. 10.1016/S0006-3495(96)79208-1

Ben-Tal, N., Honig, B., Bagdassarian, C.K., Ben-Shaul, A., 2000. Association Entropy in Adsorption Processes. Biophysical Journal 79, 1180–1187. 10.1016/S0006-3495(00)76372-7

Berendsen, H.J.C., Postma, J.P.M., Van Gunsteren, W.F., DiNola, A., Haak, J.R., 1984. Molecular dynamics with coupling to an external bath. The Journal of Chemical Physics 81, 3684–3690. 10.1063/1.448118

Bergler, H., Abraham, D., Aschauer, H., Turnowsky, F., 1994. Inhibition of lipid biosynthesis induces the expression of the pspA gene. Microbiology 140, 1937–1944. 10.1099/13500872-140-8-1937

Bertin, A., de Franceschi, N., de la Mora, E., Maity, S., Alqabandi, M., Miguet, N., di Cicco, A., Roos, W.H., Mangenot, S., Weissenhorn, W., Bassereau, P., 2020. Human ESCRT-III polymers assemble on positively curved membranes and induce helical membrane tube formation. Nat Commun 11, 2663. 10.1038/s41467-020-16368-5

Bhatia, H., Ingólfsson, H.I., Carpenter, T.S., Lightstone, F.C., Bremer, P.-T., 2019. MemSurfer: A Tool for Robust Computation and Characterization of Curved Membranes. J. Chem. Theory Comput. 15, 6411–6421. 10.1021/acs.jctc.9b00453

Bhatt, V.S., Ashley, R., Sundborger-Lunna, A., 2021. Amphipathic Motifs Regulate N-BAR Protein Endophilin B1 Auto-inhibition and Drive Membrane Remodeling. Structure 29, 61–69.e3. 10.1016/j.str.2020.09.012

Bodon, G., Chassefeyre, R., Pernet-Gallay, K., Martinelli, N., Effantin, G., Hulsik, D.L., Belly, A., Goldberg, Y., Chatellard-Causse, C., Blot, B., Schoehn, G., Weissenhorn, W., Sadoul, R., 2011. Charged Multivesicular Body Protein 2B (CHMP2B) of the Endosomal Sorting Complex Required for Transport-III (ESCRT-III) Polymerizes into Helical Structures Deforming the Plasma Membrane. Journal of Biological Chemistry 286, 40276–40286. 10.1074/jbc.M111.283671

Brissette, J.L., Russel, M., Weiner, L., Model, P., 1990. Phage shock protein, a stress protein of Escherichia coli. Proceedings of the National Academy of Sciences 87, 862–866. 10.1073/pnas.87.3.862

Buchkovich, N.J., Henne, W.M., Tang, S., Emr, S.D., 2013. Essential N-Terminal Insertion Motif Anchors the ESCRT-III Filament during MVB Vesicle Formation. Developmental Cell 27, 201–214. 10.1016/j.devcel.2013.09.009

Bureau, H.R., Merz, D.R., Hershkovits, E., Quirk, S., Hernandez, R., 2015. Constrained Unfolding of a Helical Peptide: Implicit versus Explicit Solvents. PLoS ONE 10, e0127034. 10.1371/journal.pone.0127034

Case, D.A., Aktulga, H.M., Belfon, K., Cerutti, D.S., Cisneros, G.A., Cruzeiro, V.W.D., Forouzesh, N., Giese, T.J., Götz, A.W., Gohlke, H., Izadi, S., Kasavajhala, K., Kaymak, M.C., King, E., Kurtzman, T., Lee, T.-S., Li, P., Liu, J., Luchko, T., Luo, R., Manathunga, M., Machado, M.R., Nguyen, H.M., O’Hearn, K.A., Onufriev, A.V., Pan, F., Pantano, S., Qi, R., Rahnamoun, A., Risheh, A., Schott-Verdugo, S., Shajan, A., Swails, J., Wang, J., Wei, H., Wu, X., Wu, Y., Zhang, S., Zhao, S., Zhu, Q., Cheatham, T.E., Roe, D.R., Roitberg, A., Simmerling, C., York, D.M., Nagan, M.C., Merz, K.M., 2023. AmberTools. J. Chem. Inf. Model. 63, 6183–6191. 10.1021/acs.jcim.3c01153

Cashikar, A.G., Shim, S., Roth, R., Maldazys, M.R., Heuser, J.E., Hanson, P.I., 2014. Structure of cellular ESCRT-III spirals and their relationship to HIV budding. eLife 3, e02184. 10.7554/eLife.02184

Chen, Z., Rand, R.P., 1998. Comparative Study of the Effects of Several n-Alkanes on Phospholipid Hexagonal Phases. Biophysical Journal 74, 944–952. 10.1016/S0006-3495(98)74017-2

Croll, T.I., 2018. ISOLDE: a physically realistic environment for model building into low-resolution electron-density maps. Acta Cryst D 74, 519–530. 10.1107/S2059798318002425

Cui, H., Lyman, E., Voth, G.A., 2011. Mechanism of Membrane Curvature Sensing by Amphipathic Helix Containing Proteins. Biophysical Journal 100, 1271–1279. 10.1016/j.bpj.2011.01.036

Cui, H., Mim, C., Vázquez, F.X., Lyman, E., Unger, V.M., Voth, G.A., 2013. Understanding the Role of Amphipathic Helices in N-BAR Domain Driven Membrane Remodeling. Biophysical Journal 104, 404–411. 10.1016/j.bpj.2012.12.006

Darré, L., Machado, M.R., Dans, P.D., Herrera, F.E., Pantano, S., 2010. Another Coarse Grain Model for Aqueous Solvation: WAT FOUR? J. Chem. Theory Comput. 6, 3793–3807. 10.1021/ct100379f

Darwin, A.J., 2005. The phage-shock-protein response. Molecular Microbiology 57, 621–628. 10.1111/j.1365-2958.2005.04694.x

DeLano, W., 2002. PyMOL. DeLano Scientific, San Carlos, CA 1–15.

Di Giulio, M., 2021. The phylogenetic distribution of the cell division system would not imply a cellular LUCA but a progenotic LUCA. Biosystems 210, 104563. 10.1016/j.biosystems.2021.104563

Dickson, C.J., Walker, R.C., Gould, I.R., 2022. Lipid21: Complex Lipid Membrane Simulations with AMBER. J. Chem. Theory Comput. 18, 1726–1736. 10.1021/acs.jctc.1c01217

Dolinsky, T.J., Nielsen, J.E., McCammon, J.A., Baker, N.A., 2004. PDB2PQR: an automated pipeline for the setup of Poisson-Boltzmann electrostatics calculations. Nucleic Acids Research 32, W665–W667. 10.1093/nar/gkh381

Emsley, P., Lohkamp, B., Scott, W.G., Cowtan, K., 2010. Features and development of Coot. Acta Crystallogr D Biol Crystallogr 66, 486–501. 10.1107/S0907444910007493

Finkelstein, A.V., Janin, J., 1989. The price of lost freedom: entropy of bimolecular complex formation. Protein Eng Des Sel 3, 1–3. 10.1093/protein/3.1.1

Fogolari, F., Maloku, O., Dongmo Foumthuim, C.J., Corazza, A., Esposito, G., 2018. PDB2ENTROPY and PDB2TRENT: Conformational and Translational–Rotational Entropy from Molecular Ensembles. J. Chem. Inf. Model. 58, 1319–1324. 10.1021/acs.jcim.8b00143

Ford, M.G.J., Mills, I.G., Peter, B.J., Vallis, Y., Praefcke, G.J.K., Evans, P.R., McMahon, H.T., 2002. Curvature of clathrin-coated pits driven by epsin. Nature 419, 361–366. 10.1038/nature01020

Gasteiger, E., Hoogland, C., Gattiker, A., Duvaud, S., Wilkins, M.R., Appel, R.D., Bairoch, A., Gasteiger E, Hoogland C, Gattiker A, Duvaud S, Wilkins MR, Appel RD, Bairoch A, 2005. Protein Identification and Analysis Tools on the ExPASy Server, in: The Proteomics Protocols Handbook. pp. 571–607.

Ghazi-Tabatabai, S., Saksena, S., Short, J.M., Pobbati, A.V., Veprintsev, D.B., Crowther, R.A., Emr, S.D., Egelman, E.H., Williams, R.L., 2008. Structure and Disassembly of Filaments Formed by the ESCRT-III Subunit Vps24. Structure 16, 1345–1356. 10.1016/j.str.2008.06.010

Ghosh, G., Fernández, G., 2020. pH- and concentration-dependent supramolecular self-assembly of a naturally occurring octapeptide. Beilstein J Org Chem 16, 2017–2025. 10.3762/bjoc.16.168

Goddard, T.D., Huang, C.C., Meng, E.C., Pettersen, E.F., Couch, G.S., Morris, J.H., Ferrin, T.E., 2018. UCSF ChimeraX: Meeting modern challenges in visualization and analysis. Protein Science 27, 14–25. 10.1002/pro.3235

Grossfield, A., 2024. WHAM: the weighted histogram analysis method [WWW Document]. URL http://membrane.urmc.rochester.edu/wordpress/?page_id=126

Gupta, T.K., Klumpe, S., Gries, K., Heinz, S., Wietrzynski, W., Ohnishi, N., Niemeyer, J., Spaniol, B., Schaffer, M., Rast, A., Ostermeier, M., Strauss, M., Plitzko, J.M., Baumeister, W., Rudack, T., Sakamoto, W., Nickelsen, J., Schuller, J.M., Schroda, M., Engel, B.D., 2021. Structural basis for VIPP1 oligomerization and maintenance of thylakoid membrane integrity. Cell 184, 3643–3659.e23. 10.1016/j.cell.2021.05.011

Hanson, P.I., Roth, R., Lin, Y., Heuser, J.E., 2008. Plasma membrane deformation by circular arrays of ESCRT-III protein filaments. Journal of Cell Biology 180, 389–402. 10.1083/jcb.200707031

Harris, C.R., Millman, K.J., Van Der Walt, S.J., Gommers, R., Virtanen, P., Cournapeau, D., Wieser, E., Taylor, J., Berg, S., Smith, N.J., Kern, R., Picus, M., Hoyer, S., Van Kerkwijk, M.H., Brett, M., Haldane, A., Del Río, J.F., Wiebe, M., Peterson, P., Gérard-Marchant, P., Sheppard, K., Reddy, T., Weckesser, W., Abbasi, H., Gohlke, C., Oliphant, T.E., 2020. Array programming with NumPy. Nature 585, 357–362. 10.1038/s41586-020-2649-2

Helfrich, W., 1973. Elastic Properties of Lipid Bilayers: Theory and Possible Experiments. Zeitschrift für Naturforschung C 28, 693–703. 10.1515/znc-1973-11-1209

Henne, W.M., Buchkovich, N.J., Emr, S.D., 2011. The ESCRT pathway. Dev. Cell 21, 77–91. 10.1016/j.devcel.2011.05.015

Hinshaw, J.E., Schmid, S.L., 1995. Dynamin self-assembles into rings suggesting a mechanism for coated vesicle budding. Nature 374, 190–192. 10.1038/374190a0

Hopkins, C.W., Le Grand, S., Walker, R.C., Roitberg, A.E., 2015. Long-Time-Step Molecular Dynamics through Hydrogen Mass Repartitioning. J. Chem. Theory Comput. 11, 1864–1874. 10.1021/ct5010406

Hu, M., Briguglio, J.J., Deserno, M., 2012. Determining the Gaussian Curvature Modulus of Lipid Membranes in Simulations. Biophysical Journal 102, 1403–1410. 10.1016/j.bpj.2012.02.013

Huber, S.T., Mostafavi, S., Mortensen, S.A., Sachse, C., 2020. Structure and assembly of ESCRT-III helical Vps24 filaments. Sci Adv 6, eaba4897. 10.1126/sciadv.aba4897

Humphrey, W., Dalke, A., Schulten, K., 1996. VMD: Visual molecular dynamics. Journal of Molecular Graphics 14, 33–38. 10.1016/0263-7855(96)00018-5

Hurley, J.H., Hanson, P.I., 2010. Membrane budding and scission by the ESCRT machinery: it’s all in the neck. Nat. Rev. Mol. Cell Biol. 11, 556–566. 10.1038/nrm2937

Huvet, M., Toni, T., Sheng, X., Thorne, T., Jovanovic, G., Engl, C., Buck, M., Pinney, J.W., Stumpf, M.P.H., 2011. The evolution of the phage shock protein response system: Interplay between protein function, genomic organization, and system function. Molecular Biology and Evolution 28, 1141–1155. 10.1093/molbev/msq301

Jakobi, A.J., Wilmanns, M., Sachse, C., 2017. Model-based local density sharpening of cryo-EM maps. eLife 6, e27131. 10.7554/eLife.27131

Jorgensen, W.L., Chandrasekhar, J., Madura, J.D., Impey, R.W., Klein, M.L., 1983. Comparison of simple potential functions for simulating liquid water. The Journal of Chemical Physics 79, 926–935. 10.1063/1.445869

Jovanovic, G., Mehta, P., McDonald, C., Davidson, A.C., Uzdavinys, P., Ying, L., Buck, M., 2014. The N-terminal amphipathic helices determine regulatory and effector functions of phage shock protein A (PspA) in escherichia coli. Journal of Molecular Biology 426, 1498–1511. 10.1016/j.jmb.2013.12.016

Jovanovic, G., Weiner, L., Model, P., 1996. Identification, nucleotide sequence, and characterization of PspF, the transcriptional activator of the Escherichia coli stress-induced psp operon. Journal of bacteriology 178, 1936–45. 10.1128/jb.178.7.1936-1945.1996

Junglas, B., Huber, S.T., Heidler, T., Schlösser, L., Mann, D., Hennig, R., Clarke, M., Hellmann, N., Schneider, D., Sachse, C., 2021. PspA adopts an ESCRT-III-like fold and remodels bacterial membranes. Cell 184, 3674–3688.e18. 10.1016/j.cell.2021.05.042

Junglas, B., Hudina, E., Schönnenbeck, P., Ritter, I., Heddier, A., Santiago-Schübel, B., Huesgen, P.F., Schneider, D., Sachse, C., 2024a. Structural plasticity of bacterial ESCRT-III protein PspA in higher-order assemblies. 10.1101/2024.07.08.602472

Junglas, B., Kartte, D., Kutzner, M., Hellmann, N., Ritter, I., Schneider, D., Sachse, C., 2024b. Structural basis for Vipp1 membrane binding: From loose coats and carpets to ring and rod assemblies. 10.1101/2024.07.08.602470

Junglas, B., Orru, R., Axt, A., Siebenaller, C., Steinchen, W., Heidrich, J., Hellmich, U.A., Hellmann, N., Wolf, E., Weber, S.A.L., Schneider, D., 2020. IM30 IDPs form a membrane-protective carpet upon super-complex disassembly. Communications Biology 3, 1–10. 10.1038/s42003-020-01314-4

Kleerebezem, M., Tommassen, J., 1993. Expression of the pspA gene stimulates efficient protein export in Escherichia coli. Molecular Microbiology 7, 947–956. 10.1111/j.1365-2958.1993.tb01186.x

Kobayashi, H., Yamamoto, M., Aono, R., 1998. Appearance of a stress-response protein, phage-shock protein A, in Escherichia coli exposed to hydrophobic organic solvents. Microbiology 144, 353–359. 10.1099/00221287-144-2-353

Kobayashi, R., Suzuki, T., Yoshida, M., 2007. Escherichia coli phage-shock protein A (PspA) binds to membrane phospholipids and repairs proton leakage of the damaged membranes. Molecular Microbiology 66, 100–109. 10.1111/j.1365-2958.2007.05893.x

Kozlov, M.M., McMahon, H.T., Chernomordik, L.V., 2010. Protein-driven membrane stresses in fusion and fission. Trends in Biochemical Sciences 35, 699–706. 10.1016/j.tibs.2010.06.003

Krause, D., 2019. JUWELS: Modular Tier-0/1 Supercomputer at the Jülich Supercomputing Centre. Journal of large-scale research facilities JLSRF 5, 135. 10.17815/jlsrf-5-171

Lata, S., Schoehn, G., Jain, A., Pires, R., Piehler, J., Gőttlinger, H.G., Weissenhorn, W., 2008. Helical Structures of ESCRT-III Are Disassembled by VPS4. Science 321, 1354–1357. 10.1126/science.1161070

Le Grand, S., Götz, A.W., Walker, R.C., 2013. SPFP: Speed without compromise—A mixed precision model for GPU accelerated molecular dynamics simulations. Computer Physics Communications 184, 374–380. 10.1016/j.cpc.2012.09.022

Liu, J., Tassinari, M., Souza, D.P., Naskar, S., Noel, J.K., Bohuszewicz, O., Buck, M., Williams, T.A., Baum, B., Low, H.H., 2021. Bacterial Vipp1 and PspA are members of the ancient ESCRT-III membrane-remodeling superfamily. Cell 184, 3660–3673.e18. 10.1016/j.cell.2021.05.041

Lloyd, L.J., Jones, S.E., Jovanovic, G., Gyaneshwar, P., Rolfe, M.D., Thompson, A., Hinton, J.C., Buck, M., 2004. Identification of a new member of the phage shock protein response in Escherichia coli, the phage shock protein G (PspG). Journal of Biological Chemistry 279, 55707–55714. 10.1074/jbc.M408994200

Lomize, A.L., Todd, S.C., Pogozheva, I.D., 2022. Spatial arrangement of proteins in planar and curved membranes by PPM 3.0. Protein Science 31, 209–220. 10.1002/pro.4219

Machado, M.R., Barrera, E.E., Klein, F., Sóñora, M., Silva, S., Pantano, S., 2019. The SIRAH 2.0 Force Field: Altius, Fortius, Citius. J. Chem. Theory Comput. 15, 2719–2733. 10.1021/acs.jctc.9b00006

Machado, M.R., Pantano, S., 2016. SIRAH tools: mapping, backmapping and visualization of coarse-grained models. Bioinformatics 32, 1568–1570. 10.1093/bioinformatics/btw020

Maier, J.A., Martinez, C., Kasavajhala, K., Wickstrom, L., Hauser, K.E., Simmerling, C., 2015. ff14SB: Improving the Accuracy of Protein Side Chain and Backbone Parameters from ff99SB. J. Chem. Theory Comput. 11, 3696–3713. 10.1021/acs.jctc.5b00255

Maity, S., Caillat, C., Miguet, N., Sulbaran, G., Effantin, G., Schoehn, G., Roos, W.H., Weissenhorn, W., 2019. VPS4 triggers constriction and cleavage of ESCRT-III helical filaments. Sci Adv 5, eaau7198. 10.1126/sciadv.aau7198

Manganelli, R., Gennaro, M.L., 2017. Protecting from Envelope Stress: Variations on the Phage-Shock-Protein Theme. Trends in Microbiology 25, 205–216. 10.1016/j.tim.2016.10.001

Martínez, L., Andrade, R., Birgin, E.G., Martínez, J.M., 2009. P ACKMOL : A package for building initial configurations for molecular dynamics simulations. J Comput Chem 30, 2157–2164. 10.1002/jcc.21224

McCullough, J., Clippinger, A.K., Talledge, N., Skowyra, M.L., Saunders, M.G., Naismith, T.V., Colf, L.A., Afonine, P., Arthur, C., Sundquist, W.I., Hanson, P.I., Frost, A., 2015. Structure and membrane remodeling activity of ESCRT-III helical polymers. Science 350, 1548–1551. 10.1126/science.aad8305

McDonald, C., Jovanovic, G., Ces, O., Buck, M., 2015. Membrane stored curvature elastic stress modulates recruitment of maintenance proteins pspa and vipp1. mBio 6, e01188–15. 10.1128/mBio.01188-15

McDonald, C., Jovanovic, G., Wallace, B.A., Ces, O., Buck, M., 2017. Structure and function of PspA and Vipp1 N-terminal peptides: Insights into the membrane stress sensing and mitigation. Biochimica et Biophysica Acta - Biomembranes 1859, 28–39. 10.1016/j.bbamem.2016.10.018

Megrabov, A.G., 2014. On divergence representations of the Gaussian and the mean curvature of surfaces and applications.

Melnikov, N., Junglas, B., Halbi, G., Nachmias, D., Zerbib, E., Gueta, N., Upcher, A., Zalk, R., Sachse, C., Bernheim-Groswasser, A., Elia, N., 2025. The Asgard archaeal ESCRT-III system forms helical filaments and remodels eukaryotic-like membranes. EMBO J. 10.1038/s44318-024-00346-4

Melnikov, N., Junglas, B., Halbi, G., Nachmias, D., Zerbib, E., Upcher, A., Zalk, R., Sachse, C., Bernheim-Groswasser, A., Elia, N., 2022. The Asgard archaeal ESCRT-III system forms helical filaments and remodels eukaryotic membranes, shedding light on the emergence of eukaryogenesis. 10.1101/2022.09.07.506706

Meurer, A., Smith, C.P., Paprocki, M., Čertík, O., Kirpichev, S.B., Rocklin, M., Kumar, Am., Ivanov, S., Moore, J.K., Singh, S., Rathnayake, T., Vig, S., Granger, B.E., Muller, R.P., Bonazzi, F., Gupta, H., Vats, S., Johansson, F., Pedregosa, F., Curry, M.J., Terrel, A.R., Roučka, Š., Saboo, A., Fernando, I., Kulal, S., Cimrman, R., Scopatz, A., 2017. SymPy: symbolic computing in Python. PeerJ Computer Science 3, e103. 10.7717/peerj-cs.103

Micsonai, A., Moussong, É., Wien, F., Boros, E., Vadászi, H., Murvai, N., Lee, Y.-H., Molnár, T., Réfrégiers, M., Goto, Y., Tantos, Á., Kardos, J., 2022. BeStSel: webserver for secondary structure and fold prediction for protein CD spectroscopy. Nucleic Acids Research 50, W90– W98. 10.1093/nar/gkac345

Morin, A., Eisenbraun, B., Key, J., Sanschagrin, P.C., Timony, M.A., Ottaviano, M., Sliz, P., 2013. Collaboration gets the most out of software. eLife 2, e01456. 10.7554/eLife.01456

Moser von Filseck, J., Barberi, L., Talledge, N., Johnson, I.E., Frost, A., Lenz, M., Roux, A., 2020. Anisotropic ESCRT-III architecture governs helical membrane tube formation. Nat Commun 11, 1516. 10.1038/s41467-020-15327-4

Moutoussamy, E.E., Khan, H.M., Roberts, M.F., Gershenson, A., Chipot, C., Reuter, N., 2022. Standard Binding Free Energy and Membrane Desorption Mechanism for a Phospholipase C. J. Chem. Inf. Model. 62, 6602–6613. 10.1021/acs.jcim.1c01543

Nagle, J.F., 2013. Introductory Lecture: Basic quantities in model biomembranes. Faraday Discuss. 161, 11–29. 10.1039/C2FD20121F

Nugent, T., Jones, D.T., 2013. Membrane protein orientation and refinement using a knowledge-based statistical potential. BMC Bioinformatics 14, 276. 10.1186/1471-2105-14-276

Olsson, M.H.M., Søndergaard, C.R., Rostkowski, M., Jensen, J.H., 2011. PROPKA3: Consistent Treatment of Internal and Surface Residues in Empirical p *K* _a_ Predictions. J. Chem. Theory Comput. 7, 525–537. 10.1021/ct100578z

Osadnik, H., Schöpfel, M., Heidrich, E., Mehner, D., Lilie, H., Parthier, C., Risselada, H.J., Grubmüller, H., Stubbs, M.T., Brüser, T., 2015. PspF-binding domain PspA1-144and the PspA·F complex: New insights into the coiled-coil-dependent regulation of AAA+ proteins. Molecular Microbiology 98, 743–759. 10.1111/mmi.13154

Pettersen, E.F., Goddard, T.D., Huang, C.C., Meng, E.C., Couch, G.S., Croll, T.I., Morris, J.H., Ferrin, T.E., 2021. UCSF ChimeraX: Structure visualization for researchers, educators, and developers. Protein Sci 30, 70–82. 10.1002/pro.3943

Pfitzner, A.-K., Mercier, V., Jiang, X., Moser von Filseck, J., Baum, B., Šarić, A., Roux, A., 2020. An ESCRT-III Polymerization Sequence Drives Membrane Deformation and Fission. Cell 182, 1140–1155.e18. 10.1016/j.cell.2020.07.021

Popp, P.F., Gumerov, V.M., Andrianova, E.P., Bewersdorf, L., Mascher, T., Zhulin, I.B., Wolf, D., 2022. Phyletic Distribution and Diversification of the Phage Shock Protein Stress Response System in Bacteria and Archaea. mSystems e0134821. 10.1128/msystems.01348-21

Punjani, A., Rubinstein, J.L., Fleet, D.J., Brubaker, M.A., 2017. cryoSPARC: algorithms for rapid unsupervised cryo-EM structure determination. Nature Methods 14, 290–296. 10.1038/nmeth.4169

Roe, D.R., Cheatham, T.E., 2013. PTRAJ and CPPTRAJ: Software for Processing and Analysis of Molecular Dynamics Trajectory Data. J. Chem. Theory Comput. 9, 3084–3095. 10.1021/ct400341p

Rueda-Contreras, M.D., Gallen, A.F., Romero-Arias, J.R., Hernandez-Machado, A., Barrio, R.A., 2021. On Gaussian curvature and membrane fission. Sci Rep 11, 9562. 10.1038/s41598-021-88851-y

Rueden, C.T., Schindelin, J., Hiner, M.C., DeZonia, B.E., Walter, A.E., Arena, E.T., Eliceiri, K.W., 2017. ImageJ2: ImageJ for the next generation of scientific image data. BMC Bioinformatics 18, 529. 10.1186/s12859-017-1934-z

Ryckaert, J.-P., Ciccotti, G., Berendsen, H.J.C., 1976. Numerical integration of the Cartesian Equations of Motion of a System with Constraints: Molecular Dynamics of n-Alkanes.

Schlösser, L., Sachse, C., Low, H.H., Schneider, D., 2023. Conserved structures of ESCRT-III superfamily members across domains of life. Trends in Biochemical Sciences 48, 993–1004. 10.1016/j.tibs.2023.08.009

Schmidt, O., Teis, D., 2012. The ESCRT machinery. Curr Biol 22, R116–R120. 10.1016/j.cub.2012.01.028

Schott-Verdugo, S., Gohlke, H., 2019. PACKMOL-Memgen: A Simple-To-Use, Generalized Workflow for Membrane-Protein–Lipid-Bilayer System Building. J. Chem. Inf. Model. 59, 2522–2528. 10.1021/acs.jcim.9b00269

Schrödinger, n.d. The PyMOL Molecular Graphics System.

Schroeder, W., Martin, K., Lorensen, B., 2018. The Visualization Toolkit.

Siebenaller, C., Junglas, B., Schneider, D., 2019. Functional Implications of Multiple IM30 Oligomeric States. Front. Plant Sci. 10. 10.3389/fpls.2019.01500

Smart, O.S., Goodfellow, J.M., Wallace, B.A., 1993. The pore dimensions of gramicidin A. Biophys J 65, 2455–2460.

Standar, K., Mehner, D., Osadnik, H., Berthelmann, F., Hause, G., Lünsdorf, H., Brüser, T., 2008. PspA can form large scaffolds in Escherichia coli. FEBS Letters 582, 3585–3589. 10.1016/j.febslet.2008.09.002

Steinem, C., Meinecke, M., 2021. ENTH domain-dependent membrane remodelling. Soft Matter 17, 233–240. 10.1039/C9SM02437A

Sun, C., Gonzalez, B., Jiang, W., 2022. Helical Indexing in Real Space. Sci Rep 12, 8162. 10.1038/s41598-022-11382-7

Tegunov, D., Cramer, P., 2019. Real-time cryo-electron microscopy data preprocessing with Warp. Nature Methods 16, 1146–1152. 10.1038/s41592-019-0580-y

Terwilliger, T.C., Sobolev, O.V., Afonine, P.V., Adams, P.D., 2018. Automated map sharpening by maximization of detail and connectivity. Acta Cryst D 74, 545–559. 10.1107/S2059798318004655

Thurotte, A., Schneider, D., 2019. The fusion activity of IM30 rings involves controlled unmasking of the fusogenic core. Frontiers in Plant Science 10, 108. 10.3389/fpls.2019.00108

Trott, M., 2004. The Mathematica guidebook for graphics.

Vietri, M., Schink, K.O., Campsteijn, C., Wegner, C.S., Schultz, S.W., Christ, L., Thoresen, S.B., Brech, A., Raiborg, C., Stenmark, H., 2015. Spastin and ESCRT-III coordinate mitotic spindle disassembly and nuclear envelope sealing. Nature 522, 231–235. 10.1038/nature14408

Virtanen, P., Gommers, R., Oliphant, T.E., Haberland, M., Reddy, T., Cournapeau, D., Burovski, E., Peterson, P., Weckesser, W., Bright, J., Van Der Walt, S.J., Brett, M., Wilson, J., Millman, K.J., Mayorov, N., Nelson, A.R.J., Jones, E., Kern, R., Larson, E., Carey, C.J., Polat, İ., Feng, Y., Moore, E.W., VanderPlas, J., Laxalde, D., Perktold, J., Cimrman, R., Henriksen, I., Quintero, E.A., Harris, C.R., Archibald, A.M., Ribeiro, A.H., Pedregosa, F., Van Mulbregt, P., SciPy 1.0 Contributors, Vijaykumar, A., Bardelli, A.P., Rothberg, A., Hilboll, A., Kloeckner, A., Scopatz, A., Lee, A., Rokem, A., Woods, C.N., Fulton, C., Masson, C., Häggström, C., Fitzgerald, C., Nicholson, D.A., Hagen, D.R., Pasechnik, D.V., Olivetti, E., Martin, E., Wieser, E., Silva, F., Lenders, F., Wilhelm, F., Young, G., Price, G.A., Ingold, G.-L., Allen, G.E., Lee, G.R., Audren, H., Probst, I., Dietrich, J.P., Silterra, J., Webber, J.T., Slavič, J., Nothman, J., Buchner, J., Kulick, J., Schönberger, J.L., De Miranda Cardoso, J.V., Reimer, J., Harrington, J., Rodríguez, J.L.C., Nunez-Iglesias, J., Kuczynski, J., Tritz, K., Thoma, M., Newville, M., Kümmerer, M., Bolingbroke, M., Tartre, M., Pak, M., Smith, N.J., Nowaczyk, N., Shebanov, N., Pavlyk, O., Brodtkorb, P.A., Lee, P., McGibbon, R.T., Feldbauer, R., Lewis, S., Tygier, S., Sievert, S., Vigna, S., Peterson, S., More, S., Pudlik, T., Oshima, T., Pingel, T.J., Robitaille, T.P., Spura, T., Jones, T.R., Cera, T., Leslie, T., Zito, T., Krauss, T., Upadhyay, U., Halchenko, Y.O., Vázquez-Baeza, Y., 2020. SciPy 1.0: fundamental algorithms for scientific computing in Python. Nat Methods 17, 261–272. 10.1038/s41592-019-0686-2

Williams, C.J., Headd, J.J., Moriarty, N.W., Prisant, M.G., Videau, L.L., Deis, L.N., Verma, V., Keedy, D.A., Hintze, B.J., Chen, V.B., Jain, S., Lewis, S.M., Arendall III, W.B., Snoeyink, J., Adams, P.D., Lovell, S.C., Richardson, J.S., Richardson, D.C., 2018. MolProbity: More and better reference data for improved all-atom structure validation. Protein Science 27, 293–315. 10.1002/pro.3330

Wolf, D., Kalamorz, F., Wecke, T., Juszczak, A., Mäder, U., Homuth, G., Jordan, S., Kirstein, J., Hoppert, M., Voigt, B., Hecker, M., Mascher, T., 2010. In-depth profiling of the LiaR response of bacillus subtilis. Journal of Bacteriology 192, 4680–4693. 10.1128/JB.00543-10

Zhang, X., 2022. Python-based Helix Indexer: A graphical user interface program for finding symmetry of helical assembly through Fourier-Bessel indexing of electron microscopic data. Protein Sci 31, 107–117. 10.1002/pro.4186

Zheng, S., Wolff, G., Greenan, G., Chen, Z., Faas, F.G.A., Bárcena, M., Koster, A.J., Cheng, Y., Agard, D.A., 2022. AreTomo: An integrated software package for automated marker-free, motion-corrected cryo-electron tomographic alignment and reconstruction. Journal of Structural Biology: X 6, 100068. 10.1016/j.yjsbx.2022.100068

Zhu, L., Jorgensen, J.R., Li, M., Chuang, Y.-S., Emr, S.D., 2017. ESCRTs function directly on the lysosome membrane to downregulate ubiquitinated lysosomal membrane proteins. eLife 6, e26403. 10.7554/eLife.26403

